# Letting go to hold on: loss of CCM2 disrupts endothelial force balance in cerebral cavernous malformations

**DOI:** 10.64898/2026.10.09.757835

**Authors:** Sarah Vorsselmans, Samet Aytekin, Christiane Oddou, Thomas Manicom Ramsamy, Eva Faurobert, Susana Rocha

## Abstract

Cerebral cavernous malformations (CCM) are vascular lesions characterized by impaired endothelial integrity and altered cell-cell and cell-matrix interactions. While endothelial contractility and tractions have been implicated in CCM, how these forces are transmitted and coordinated across adhesion systems remains unknown. Here, we combined genetically encoded FRET-based molecular tension sensors of vinculin and VE-cadherin with quantitative analysis of adhesion organization to dissect this force balance in CCM2-deficient endothelial monolayers. We found that CCM2 loss redistributes molecular forces, partially through ROCK-dependent contractility, across endothelial adhesions, shifting mechanical engagement from cell-cell junctions toward the cell-matrix interface. Matrix stiffness further amplifies this imbalance, as healthy endothelial cells maintain the balance between cell-cell and cell-matrix adhesions by reinforcing both in response to stiffening, whereas CCM2-deficient cells preferentially reinforce the cell-matrix interface. These findings suggest that CCM-associated matrix remodeling could reinforce an already altered mechanical state, establishing a potential feed-forward interaction between endothelial force transmission and the extracellular environment. More broadly, our results identify the CCM complex as a molecular hub controlling force balance in response to the cell mechanical environment and establish a combined tension sensing and morphology-based imaging platform for studying adhesion mechanobiology.

## Introduction

Cerebral cavernous malformations (CCM) are vascular lesions of the central nervous system characterized by clusters of dilated, thin-walled capillaries with impaired barrier function. Patients can present with seizures, focal neurological deficits and intracerebral hemorrhage (Snellings et al., 2021; Wong et al., 2000). CCM affects an estimated 0.5% of the general population and occurs in both familial and sporadic forms (Goldstein & Solomon, 2017). Familial CCM arises from inherited loss-of-function mutations in one of three genes: KRIT1/CCM1 (Laberge-le Couteulx et al., 1999), CCM2 (Denier et al., 2004; Liquori et al., 2003) or PDCD10/CCM3 (Bergametti et al., 2005). The encoded proteins assemble into a trimeric CCM complex, where CCM2 occupies a central position, bridging CCM1 and CCM3 and coordinating downstream signaling (Swamy & Glading, 2022; Voss et al., 2007). Despite advances in understanding CCM pathogenesis, treatment options are limited to surgical resection or symptom management for symptomatic lesions (Dulamea & Lupescu, 2024).

The CCM complex is increasingly recognized as an important regulator of endothelial mechanosignaling, affecting how endothelial cells generate, transmit and respond to force (Rödel & Abdelilah-Seyfried, 2026). Our understanding of the consequences of CCM loss has been pieced together from studies using distinct experimental systems and readouts. Initial work showed that CCM deficiency destabilizes cell-cell junctions and increases endothelial permeability (Glading et al., 2007; Stockton et al., 2010; Whitehead et al., 2009). These changes have been linked to altered Rap1 and SMURF1 signaling and increased RhoA activity, leading to enhanced actomyosin contractility and reorganization of the actin cytoskeleton (Borikova et al., 2010; Crose et al., 2009; Glading et al., 2007; Stockton et al., 2010; Whitehead et al., 2009). Later work extended this mechanical phenotype to adhesions at the cell and extracellular matrix (ECM) interface. CCM depletion increases β1-integrin activity and Rho-associated coiled-coil-containing kinase (ROCK) dependent contractility, reinforcing cell-ECM adhesion and promoting ECM remodeling that may further compromise cell-cell adhesion (Faurobert et al., 2013; Jilkova et al., 2014; Lisowska et al., 2018; Vannier et al., 2021). More recently, this mechanical dysregulation has been linked to disease progression in which increased forces exerted on the three-dimensional matrix contribute to the invasive phenotype of CCM-mutant cells, while force transmission through cell-cell junctions contributes to the recruitment of neighboring wild-type endothelial cells into the growing mosaic lesions (Shapeti et al., 2024). Together, these studies establish that both cell-cell and cell-ECM adhesions are mechanically dysregulated following CCM loss and that these changes are involved in lesion invasion and expansion. However, previous studies have measured forces at cellular or substrate level, leaving unresolved how CCM loss alters the forces experienced by individual adhesion molecules. Importantly, cell-cell and cell-ECM adhesions have been examined separately, making it unclear whether CCM loss simply increases endothelial force generation or disrupts how forces are distributed and coordinated between the two adhesion systems. Distinguishing between these possibilities requires measuring molecular tension and adhesion organization at both interfaces within the same endothelial system.

Genetically encoded Förster resonance energy transfer (FRET)-based molecular tension sensors provide the molecular resolution needed to achieve this. By reporting piconewton-scale forces across individual mechanosensitive proteins, these sensors provide a direct molecular readout of adhesion loading, complementing traction-based measurements (Grashoff et al., 2010). Tension sensors for vinculin (Grashoff et al., 2010) and VE-cadherin (Conway et al., 2013), key mechanosensitive components of focal adhesions (FA) and adherens junctions (AJ) respectively, enable direct comparison of cell-ECM and cell-cell force transmission within the same experimental system.

Here, we combine fluorescence lifetime imaging microscopy (FLIM)-FRET imaging of vinculin and VE-cadherin tension sensors with quantitative analysis of adhesion morphology to investigate the molecular mechanics of FAs and AJs in a CCM2-depleted endothelial model. By adjusting ROCK-dependent contractility and the mechanical properties of the extracellular environment, we examine how this altered mechanical state responds to intracellular and extracellular cues. Our findings are consistent with the CCM complex acting as a regulator of the force balance between cell-cell and cell-ECM adhesions and of how this balance is renegotiated in response to the mechanical microenvironment. They further reveal matrix stiffness as a potential amplifier of the force imbalance induced by CCM2 loss, supporting a mechanically primed state that may contribute to disease progression. Together, this integrated approach provides a molecular-scale view of the mechanical dysregulation associated with CCM loss and a framework to investigate altered mechanotransduction in disease contexts.

## Results

### An integrated imaging platform to quantify adhesion mechanics in an *in vitro* CCM model

To investigate how CCM loss alters the molecular tension and adhesion organization of cell-cell and cell-ECM adhesions, we established a FLIM-FRET imaging platform in a well-characterized *in vitro* model of CCM disease (Figure 1).

**Figure 1.**
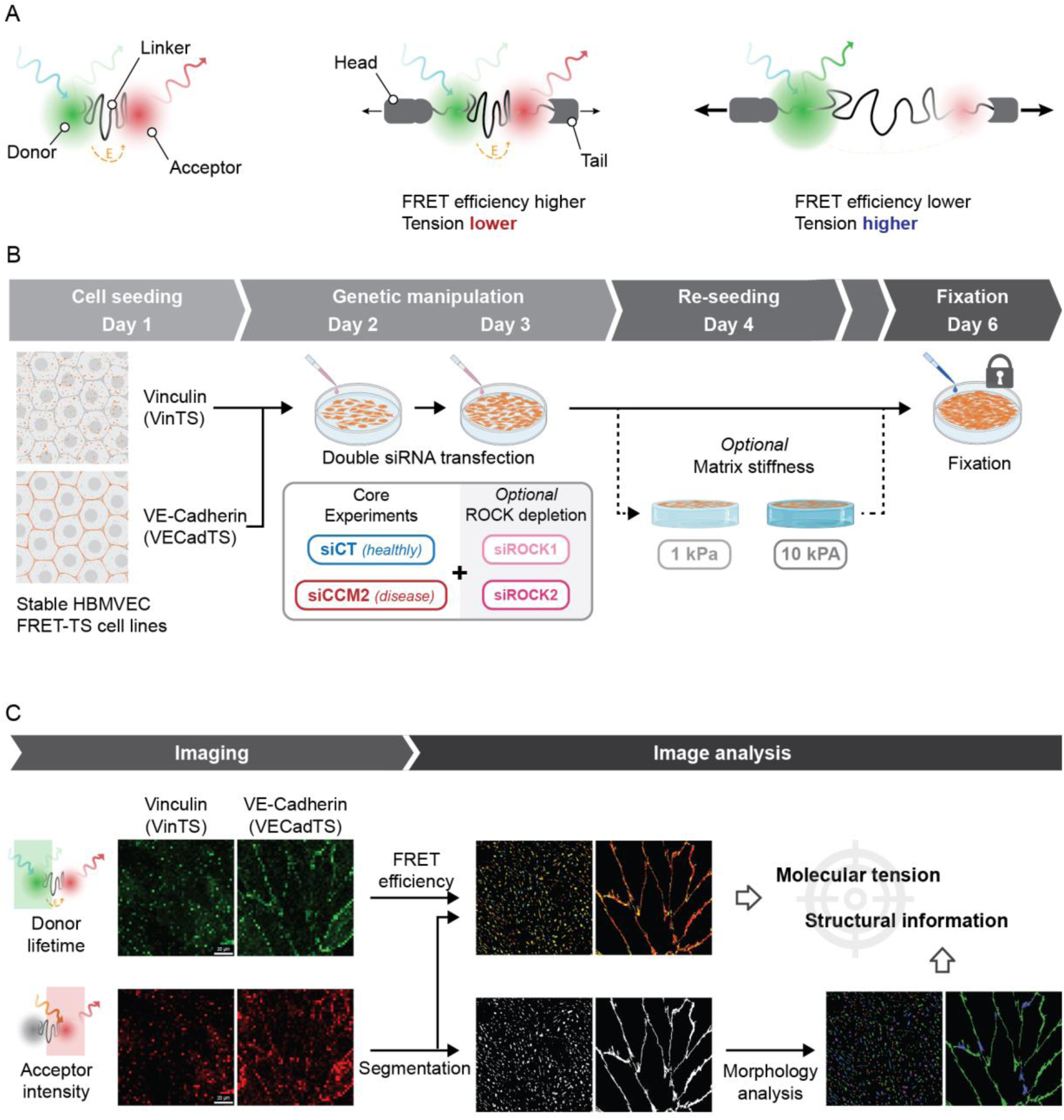
Integrated FLIM-FRET and structural imaging workflow for endothelial adhesion mechanics. **(A)** Left: Tension sensing module (TSMod) with donor, spring/elastic linker and acceptor. Right: Full tension sensor between the head and tail domain of either vinculin or VE-cadherin. When force is applied to the tension sensor, the distance between donor and acceptor increases, leading to a decrease in FRET efficiency. **(B)** Sample preparation started by seeding stable HBMVECs expressing either vinculin tension sensor (VinTS) or VE-cadherin tension sensor (VECadTS). Next, double siRNA transfection on 2 consecutive days resulted in CCM2 depletion with optional additional ROCK1 or ROCK2 depletion. Cells were reseeded on day 4 for experiments on collagen-coated PAA hydrogels of either 1 or 10 kPa. Fixation was performed on day 6. **(C)** During imaging, the FRET efficiency was obtained by imaging the donor lifetime, while intensity signal for morphology analysis was obtained from the non-FRET acceptor intensity. The full analysis pipeline performed segmentation, FRET-FLIM-phasor calculation and morphology analysis to extract molecular tension and structural information.

Endothelial forces at the cell-ECM and cell-cell interfaces were measured using genetically encoded molecular tension sensors targeted to the mechanosensitive proteins vinculin and VE-cadherin, which localize at FA and AJ, respectively (Conway et al., 2013; Grashoff et al., 2010). Both sensors consist of a tension sensing module (TSMod), genetically inserted between the head and tail domain of either vinculin or VE-cadherin (Figure 1A). The tension sensing module consists of a donor and an acceptor fluorophore capable of FRET, connected by a flexible polypeptide linker that extends under mechanical force. Forces transmitted through the protein stretch the linker, increasing the distance between the fluorophores and reducing FRET efficiency. The sensor therefore reports the molecular tension experienced by the protein, with lower FRET efficiencies indicating higher tension across the protein.

The modular design of the TSMod allows both the fluorophore pair and the elastic linker to be selected according to the requirements of the application. For the fluorophores, we selected the Clover-mScarlet-I fluorophore pair, which we recently identified as a robust FRET pair with improved dynamic range for FLIM-based molecular tension measurements (Aytekin et al., 2026). The choice of the elastic linker is particularly important because the tension range of the molecular tension sensors depends on both the mechanical properties of the linker and the protein and cell type in which the sensor operates. We therefore evaluated three previously reported linker modules that operate at different tension ranges (F40 (1-6 pN), FL (3-5 pN) and HP35st (8-12 pN)) in both vinculin and VE-cadherin sensors (Figure S1) (Austen et al., 2015; Grashoff et al., 2010; Ringer et al., 2017). Sensor performance was assessed by comparing the FRET efficiency of each full-length construct with that of its corresponding tailless (TL) variant. In the TL control, removal of the force-transmitting C-terminal tail prevents mechanical loading of the sensor, thereby providing a tension-insensitive reference for the unloaded state. Among the linkers tested, FL consistently yielded the largest difference in FRET efficiency between the tension-sensitive and VinTL constructs, and performed comparably to F40 for VE-cadherin. We therefore selected the FL linker for both sensors to maintain a consistent sensor platform across all experiments.

These selected tension sensors for vinculin (VinTS) and VE-cadherin (VECadTS) were then stably expressed in primary human brain microvascular endothelial cells (HBMVECs) (Figure 1B). To model CCM, cells were subjected to two rounds of small interfering RNA (siRNA)-mediated CCM2 silencing. Since CCM2 is an essential component of the CCM1/2 complex, its depletion also results in the dysregulation of CCM1 protein (Faurobert et al., 2013; Faurobert & Albiges-Rizo, 2010; Swamy & Glading, 2022). Beyond modeling CCM through CCM2 depletion, the platform enables the systematic interrogation of additional mechanical regulators. In this work, ROCK1 or ROCK2 was additionally silenced to modulate actomyosin contractility, while the influence of substrate mechanics was investigated by culturing cells on collagen-coated polyacrylamide (PAA) hydrogels with elastic moduli of 1 and 10 kPa. The 1 and 10 kPa substrates were selected to represent healthy brain tissue and the estimated stiffer microenvironment associated with CCM lesions, respectively (Budday et al., 2015; Frischer et al., 2008; Gautam et al., 2026; Ren et al., 2025; Urbanski et al., 2019). Efficient knockdown of all target genes was confirmed by quantitative PCR (Figure S2).

After fixation, cells were imaged by FLIM (Figure 1C). The fluorescence lifetime of the donor fluorophore was used to calculate the apparent FRET efficiency via the FLIM-phasor approach (Coucke et al., 2023; 2024; Aytekin et al., 2026), providing a readout of molecular tension. For simplicity, apparent FRET efficiency is hereafter referred to as FRET efficiency. In parallel, a sequentially acquired acceptor-channel image was used for quantitative image analysis of FAs and AJs, providing complementary information on their morphology and structural organization. Together, these measurements provide an integrated analysis of molecular tension and structural organization at both FAs and AJs within this *in vitro* CCM model.

### CCM loss remodels adhesion architecture and redistributes molecular tension from cell-cell to cell-ECM adhesions

Previous studies from our group and others have shown that CCM-deficient endothelial cells invert their adhesive balance, progressively weakening intercellular junctions while strengthening interactions with the ECM (Faurobert et al., 2013; Glading et al., 2007; Lisowska et al., 2018; Stockton et al., 2010; Whitehead et al., 2009). These observations propose a reorganization of force transmission within the endothelium, yet the distribution of molecular tension across these adhesion systems has not been investigated directly. We therefore combined molecular tension with quantitative analysis of adhesion organization to evaluate how CCM loss affects endothelial adhesion mechanics and structure.

Given the reported increase in cell-matrix interactions following CCM loss, we first investigated how CCM2 depletion influences FA organization and the molecular tension across vinculin. Vinculin tension sensors were expressed in HBMVECs under control (siCT) and CCM2-depleted (siCCM2) conditions (Figure 2A). CCM2 depletion resulted in an increase in the number of FAs (Figure 2B), consistent with previous observations in isolated CCM-silenced human umbilical vein endothelial cells (HUVECs) (Faurobert et al., 2013; Lisowska et al., 2018). FAs also showed increased alignment throughout the field of view (FOV) (Figure 2C), indicating reorganization of the adhesive network.

**Figure 2.**
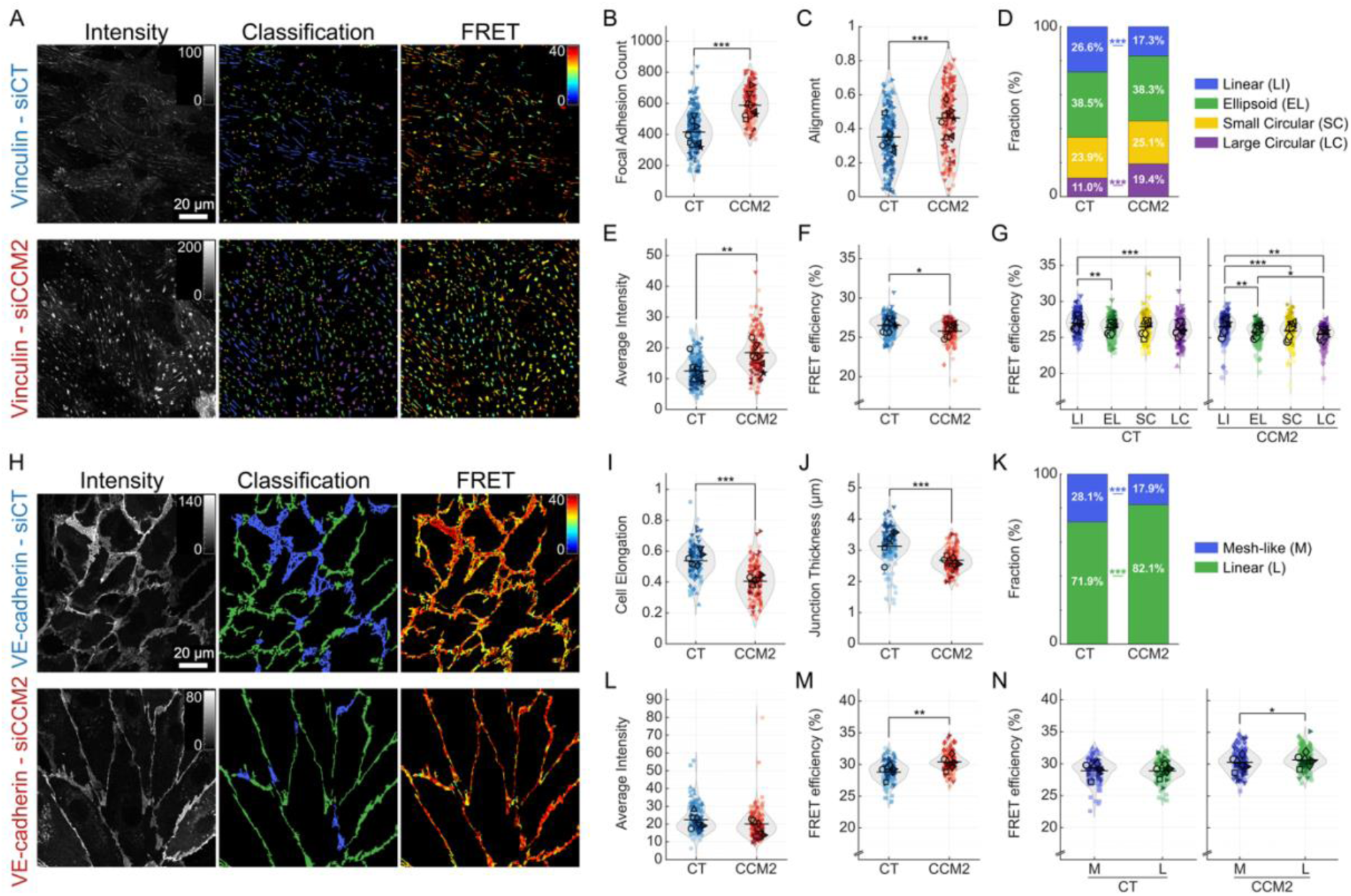
FA and AJ organization and molecular tension following CCM2 depletion. **(A-G)** Analysis of FAs using vinculin tension sensor (VinTS) **(A)** Representative images of HBMVEC monolayers stably expressing VinTS, grouped by control (siCT) and siCCM2-depleted samples. Donor fluorescence intensity images on the left, morphology classification of linear (blue), ellipsoid (green), small circular (yellow), large circular (purple) FAs in the middle panels and FRET efficiency images on the right. **(B)** FA count per FOV. **(C)** FA alignment within a given FOV (non-circular FAs only). **(D)** Stacked histogram representing the fraction of each morphological group per condition. **(E)** Average vinculin intensity representing vinculin recruitment. **(F)** Average FRET efficiency indicative of tension load. **(G)** Average FRET efficiency separated per morphological category, for CT and CCM2-depleted conditions, using the same color legend as in the panel above. (N = 217, 183 for CT and CCM2 respectively, across 8 biological replicates). Each data point represents an average value per FOV which includes multiple cells. **(H-N)** Analysis of AJs using VE-cadherin tension sensor (VECadTS) **(H)** Representative images of HBMVEC monolayers stably expressing VECadTS, grouped by control (siCT) and siCCM2-depleted samples. Donor fluorescence intensity images on the left, morphology classification of mesh-like (blue) and linear (green) AJs in the middle panels and FRET efficiency images on the right. **(I)** Average cell elongation calculated through the major and minor axis ratio of every cell. **(J)** Average AJ thickness. **(K)** Stacked histogram representing the fraction of each morphological group per condition. **(L)** Average VE-cadherin intensity representing VE-cadherin recruitment. **(M)** Average FRET efficiency indicative of tension load. **(N)** Average FRET efficiency separated per morphological category for CT and CCM2-depleted conditions, using the same color legend as in the panel above. (N = 140, 154 for CT and CCM2 respectively, across 6 biological replicates). Each data point represents an average value per FOV which includes multiple cells. All violin scatterplots are superplots distinguishing between technical replicates of each biological replicate. Statistical analysis between all groups was performed using linear-mixed effects model on the whole dataset, with only the plots per category (G, N) using rmANOVA (see Methods). *p < 0.05, **p < 0.01, ***p < 0.001.

To characterize adhesion morphology in more detail, FAs were classified into four morphological categories based on their shape and size: linear, ellipsoid, small circular and large circular adhesions (Figure 2A) (see Methods). Linear, ellipsoid and circular adhesions were distinguished based on elongation and circular FAs were further separated by area to distinguish smaller structures from the larger, force-matured adhesions (Kanchanawong & Calderwood, 2022; Tan et al., 2003). CCM2 depletion shifted the original distribution of these populations, increasing the proportion of large circular adhesions while reducing the fraction of linear adhesions (Figure 2D and Figure S3A).

We also assessed vinculin recruitment by measuring fluorescence intensity as a proxy for local vinculin density. Our analysis showed that CCM2 depletion increased vinculin density within FAs (Figure 2E). This effect was observed across all morphological categories (Figure S3B). Interestingly, large circular FAs displayed the highest vinculin density in both control and CCM2-depleted conditions (Figure S4), indicating preferential vinculin accumulation within these structures.

In addition to these changes in FA organization and vinculin recruitment, CCM2 depletion also altered the mechanical loading of vinculin. FRET efficiency was lower in CCM2-depleted cells, corresponding to increased tension across vinculin molecules (Figure 2F). In contrast, no difference was observed with the tension-insensitive control (Figure S5), confirming that changes in FRET reflected mechanical loading of vinculin rather than an effect of CCM2 silencing on sensor behavior. To determine whether this increase was associated with a particular FA morphology, the analysis was repeated for each adhesion category separately. All adhesion classes displayed the same trend toward lower FRET efficiency following CCM2 depletion (Figure S3C), indicating that the increase in vinculin tension was not restricted to a specific FA morphology. However, the magnitude of vinculin tension changed between FA categories in both control and CCM2-depleted cells. Large circular FAs consistently exhibited the lowest FRET efficiency, whereas linear adhesions displayed the highest (Figure 2G), indicating that large circular adhesions experience the highest molecular tension, independent of CCM status. Consistently, actin staining revealed that the large circular adhesions were associated with prominent stress fiber networks (Figure S6). Together, these findings indicate that CCM2 loss alters both FA organization and molecular tension, with increased vinculin recruitment and loading at cell-ECM adhesions.

To assess whether the CCM-induced shift in adhesive balance extended to cell-cell adhesions, we analyzed AJ organization and VE-cadherin tension (Figure 2H). The visualization of VE-cadherin allowed us to study cell morphology and confirm the previously described elongation of CCM2-depleted cells (Jilkova et al., 2014) (Figure 2I) and increased cell size (Figure S7). The elongated morphology was consistent with the increased FA alignment observed in CCM2-depleted cells (Figure 2C). When examining the AJs themselves, the CCM2-depleted cells displayed thinner junctions (Figure 2J and Figure S8), in line with earlier findings (Lisowska et al., 2018; Whitehead et al., 2009).

To further characterize this phenotype, we classified AJs into two morphological categories: thin linear adhesions and broad mesh-like regions, reminiscent of reticular junctions (Figure 2H) (Brandon et al., 2024; Fernández-Martín et al., 2012; Postma et al., 2025; Urbano et al., 2017). Both control and CCM2-silenced HBMVEC monolayers contained linear and mesh-like VE-cadherin regions, but CCM2 depletion increased the proportion of thin linear regions (Figure 2K and Figure S9A). In contrast with the increase in vinculin density observed in FA (Figure 2E), overall VE-cadherin density at AJs remained unchanged following CCM2 depletion (Figure 2L and Figure S9B). Analysis of the individual junctional morphologies, however, revealed differences that were not apparent from the overall measurement. In control cells, mesh-like regions displayed higher VE-cadherin density than linear AJs. Following CCM2 depletion, VE-cadherin density decreased in mesh-like regions but remained unchanged in linear AJs, such that the difference between the two junctional morphologies was no longer observed (Figure S10).

Given the observed morphological changes at the junctions, we tested whether CCM2 depletion also altered mechanical loading of VE-cadherin. FRET efficiency increased in CCM2-silenced endothelial cells, corresponding to lower tension (Figure 2M). No change was observed in the tension-insensitive control sensor (Figure S11). The decrease in tension was similar across categories (Figure S9C). When mesh-like and linear junctional regions were analyzed per condition, linear and mesh-like AJs displayed similar FRET efficiencies in control cells (Figure 2N). Following CCM2 depletion, FRET efficiency increased in both junctional morphologies. This increase was more prominent in linear AJs, resulting in higher FRET and thus lower VE-cadherin tension in linear compared with mesh-like regions (Figure 2N). These findings indicate that CCM2 depletion is associated with thinner AJs and reduced molecular tension across VE-cadherin molecules.

Taken together, these results show that CCM2 loss alters both the structure and molecular mechanics of endothelial adhesions. While FAs become more abundant, enriched in vinculin, and carry increased vinculin tension, AJs become thinner and less mesh-like, with reduced tension across VE-cadherin. These opposing responses suggest a redistribution of force transmission away from the cell-cell interface and toward the cell-ECM interface, disrupting the mechanical balance between the two adhesive systems.

### ROCK signaling regulates endothelial adhesion organization and molecular tension

Given the established roles of ROCK1 and ROCK2 in CCM pathogenesis (Ayata et al., 2024; Borikova et al., 2010; Lisowska et al., 2018; Vannier et al., 2021) and actomyosin regulation, we investigated their contribution to the altered adhesion organization and molecular tension associated with CCM2 loss. We examined how ROCK1 or ROCK2 depletion affects FAs and AJs at both the structural and mechanical levels (Figure 3A and 3G).

**Figure 3.**
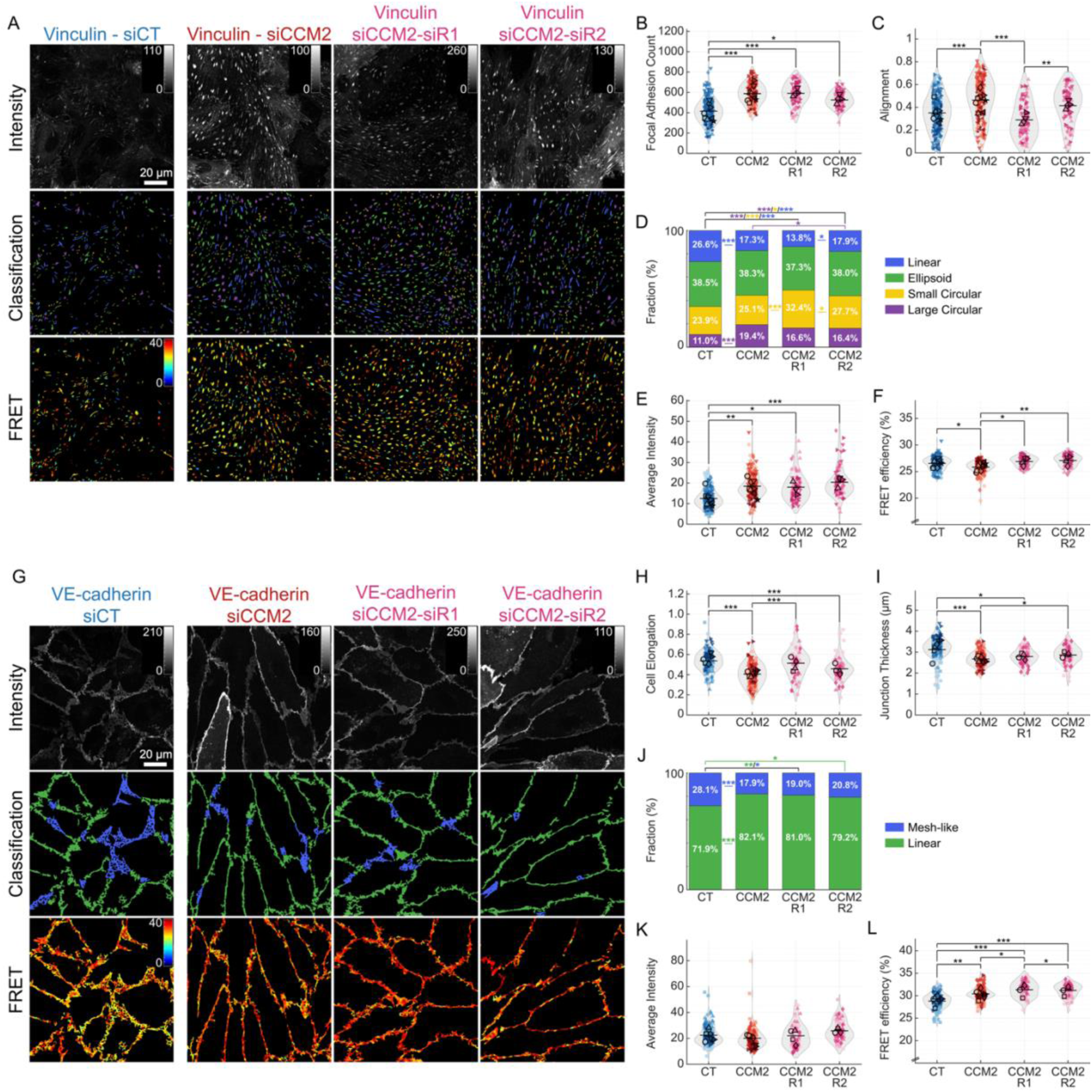
Effect of ROCK silencing on FA and AJ in CCM2-depleted monolayers. **(A-F)** Analysis of FAs using the vinculin tension sensor (VinTS) **(A)** Representative images of HBMVEC monolayers stably expressing VinTS, grouped by control (siCT), siCCM2, siCCM2-siROCK1, siCCM2-siROCK2 depleted samples. Donor fluorescence intensity images on the top, morphology classification of linear (blue), ellipsoid (green), small circular (yellow), large circular (purple) FAs in the middle panels and FRET efficiency images on the bottom. **(B)** FA count per FOV. **(C)** FA alignment within a given FOV (non-circular FAs only). **(D)** Stacked histogram representing the fraction of each morphological group per condition. **(E)** Average vinculin intensity representing vinculin recruitment. **(F)** Average FRET efficiency indicative of tension load. (N = 217, 183, 73, 73 for CT, CCM2, CCM2-ROCK1, CCM2-ROCK2 respectively, across 8 or 4 biological replicates). Each data point represents an average value per FOV which includes multiple cells. **(G-L)** Analysis of AJs using the VE-cadherin tension sensor (VECadTS) **(G)** Representative images of HBMVEC monolayers stably expressing VECadTS, grouped by control (siCT), siCCM2, siCCM2-siROCK1, siCCM2-siROCK2 depleted samples. Donor fluorescence intensity images on the top, morphology classification of mesh-like (blue) and linear (green) AJs in the middle panels and FRET efficiency images on the bottom. **(H)** Average cell elongation calculated through the major and minor axis ratio of every cell. **(I)** Average AJ thickness. **(J)** Stacked histogram representing the fraction of each morphological group per condition. **(K)** Average VE-cadherin intensity representing VE-cadherin recruitment. **(L)** Average FRET efficiency indicative of tension load. (N = 140, 154, 73, 72 for siCT, siCCM2, siCCM2-siROCK1, siCCM2-siROCK2 respectively, across 6 or 4 biological replicates). Each data point represents an average value per FOV which includes multiple cells. All violin scatterplots are superplots distinguishing between technical replicates of each biological replicate. Statistical analysis between all groups was performed using linear-mixed effects model on the whole dataset (see Methods). *p < 0.05, **p < 0.01, ***p < 0.001.

Co-silencing of CCM2 with ROCK1 revealed partial and isoform-specific effects on FA architecture in CCM2-depleted cells. Additional ROCK1 depletion did not alter overall FA abundance (Figure 3B) but significantly reduced FA alignment (Figure 3C). Fluorescence images of stained F-actin showed that this reduced alignment was accompanied by a loss of transverse stress fibers (Figure S12). Morphological analysis of the FAs further revealed a decrease in the proportion of linear and large circular adhesions, alongside an increase in small circular adhesions (Figure 3D; Figure S13A), with similar trends observed following ROCK1 depletion in control cells (Figure S14).

In contrast, CCM2-ROCK2 co-silencing had a more limited effect on FA organization. Neither FA abundance nor alignment was significantly altered relative to CCM2-silenced cells (Figure 3B-C), and transverse stress fibers remained present (Figure S12). Morphological analysis showed a selective reduction in the proportion of large circular adhesions (Figure 3D; Figure S13A), with no significant changes in the other adhesion categories. Notably, this response differed from what was observed in control cells, where ROCK2 depletion reduced FA alignment without significantly altering adhesion morphology (Figure S14).

Although ROCK1 and ROCK2 depletion had distinct effects on FA architecture, their effects on molecular loading were similar. Co-silencing of either ROCK isoform did not alter vinculin density within FAs (Figure 3E), but both reduced vinculin tension relative to CCM2-silenced cells (Figure 3F). These effects were consistently observed across the different FA categories (Figure S13B-C). Consistent with our observations in HBMVEC monolayers without additional ROCK depletion, large circular adhesions displayed the highest vinculin density across all conditions, while linear adhesions generally exhibited the lowest tension (Figure S15). However, differences in tension between adhesion categories became less pronounced following ROCK depletion (Figure S15). In control cells, ROCK1 or ROCK2 depletion had no effect on vinculin density and decreased vinculin tension, although this reduction was not significant for ROCK2 silencing (Figure S14). Thus, while ROCK1 and ROCK2 have distinct effects on FA architecture, both isoforms contribute to vinculin tension with minimal effects on vinculin recruitment.

Regarding cell shape, co-silencing of CCM2 with ROCK1 partially restored cell morphology toward control levels, resulting in a rounder cell shape compared to CCM2-silenced cells (Figure 3H). This change was consistent with the reduced FA alignment observed under the same condition (Figure 3C). Cell size remained unchanged (Figure S16). The thickness of AJs was largely maintained across conditions, although a modest increase was observed following CCM2-ROCK2 co-silencing (Figure 3I). However, neither additional ROCK1 nor ROCK2 depletion significantly altered the relative abundance of linear and mesh-like junctional regions in CCM2-silenced cells (Figure 3J; Figure S17A). In control endothelial cells, silencing of either ROCK1 or ROCK2 alone did not significantly affect cell shape or junction thickness (Figure S18).

VE-cadherin density at AJs remained largely unchanged following CCM2-ROCK1 or CCM2-ROCK2 co-silencing (Figure 3K; Figure S17B), whereas molecular tension across VE-cadherin was substantially reduced under both conditions (Figure 3L). Similar effects were observed upon ROCK depletion in control cells (Figure S18). Analysis of the individual junctional morphologies showed that VE-cadherin tension was reduced in both mesh-like and linear junctional regions (Figure S17C). Across all conditions, VE-cadherin density was similar between the two junction morphologies (Figure S19A), while linear regions consistently exhibited lower tension than mesh-like regions (Figure S19B).

Overall, ROCK depletion had relatively modest effects on the morphological phenotype induced by CCM2 loss in HBMVECs, although ROCK1 had a more pronounced effect on cellular and FA organization than ROCK2. Despite these distinct structural responses, both ROCK isoforms reduced molecular tension across vinculin and VE-cadherin. These effects were more consistent than the structural changes, indicating that ROCK-dependent contractility contributes to force transmission at both cell-ECM and cell-cell interfaces in CCM2-deficient endothelial cells.

### CCM2 loss disrupts coordinated mechanoadaptation to matrix stiffness

ECM remodeling has been observed in human CCM lesions (Clatterbuck et al., 2001; Kiliç et al., 2000; Rothbart et al., 1996), CCM mouse models (Faurobert et al., 2013; Hong et al., 2020), and *in vitro* models (Faurobert et al., 2013; Shapeti et al., 2024; Vannier et al., 2021), suggesting that endothelial cells may encounter an altered mechanical microenvironment during disease progression. We therefore investigated whether CCM2-deficient endothelial cells respond differently to matrix stiffness.

HBMVECs were cultured on collagen-coated PAA gels of 1 kPa and 10 kPa (Figure 4A,G). A stiffness of 1 kPa approximates indentation-based values for healthy brain tissue (Budday et al., 2015), whereas 10 kPa was selected as an estimate of a mechanically altered CCM microenvironment based on the collagen-rich, hyalinized, and calcified histopathology of CCM lesions (Frischer et al., 2008; Ren et al., 2025) and mechanical measurements from other fibrotic CNS pathologies (Gautam et al., 2026; Urbanski et al., 2019). As no direct mechanical measurements of CCM lesions have been reported, 10 kPa should be considered a literature-based estimate rather than a measured lesion stiffness.

**Figure 4.**
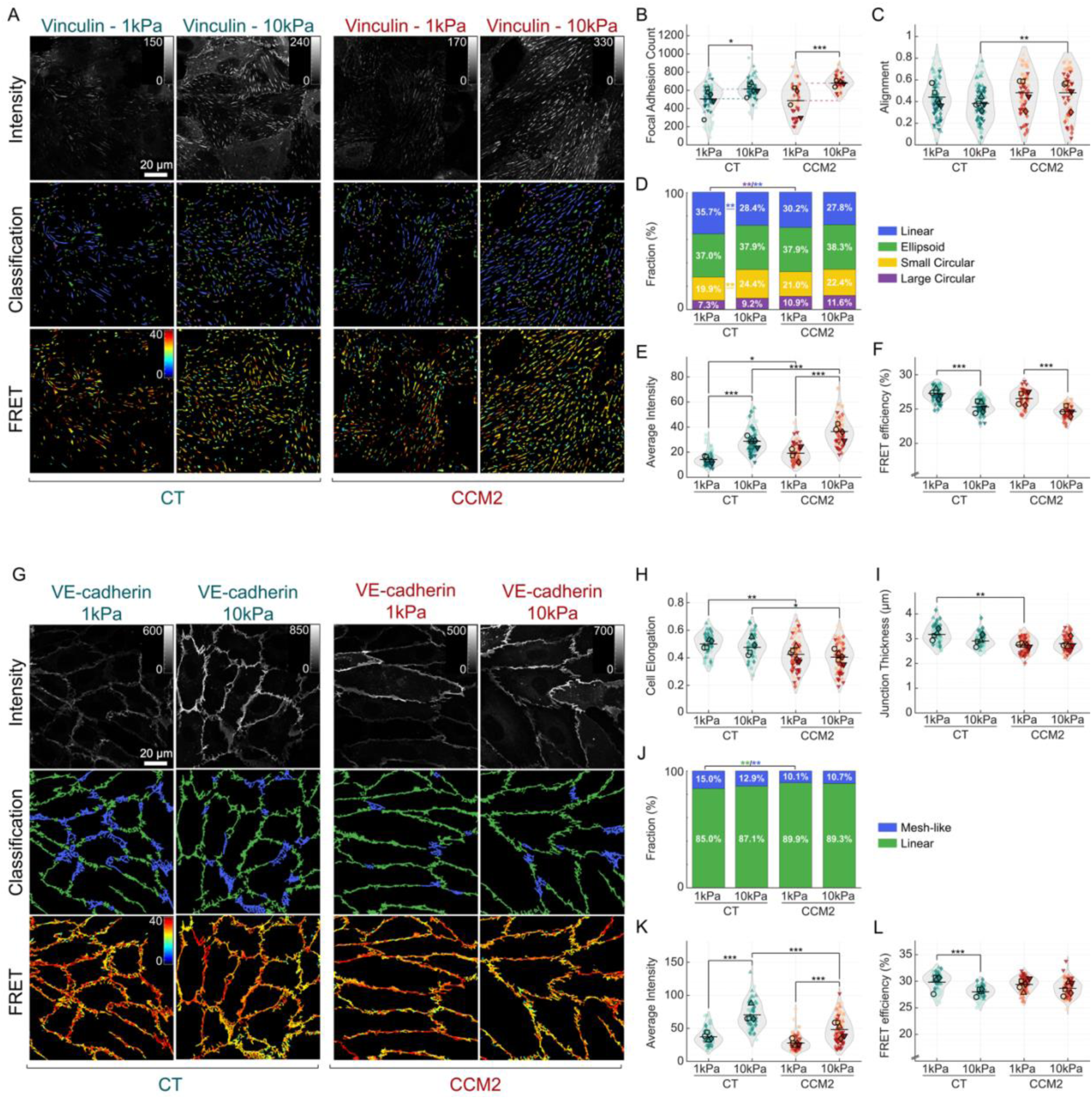
Increased stiffness exacerbates unbalanced endothelial cell adhesions in CCM2-depleted HBMVECs. **(A-F)** Analysis of FAs using the vinculin tension sensor (VinTS) **(A)** Representative images of HBMVEC monolayers stably expressing VinTS, on 1 and 10 kPa collagen-coated PAA gels grouped by control (siCT) and siCCM2-depleted samples. Donor fluorescence intensity images in the top row, morphology classification of linear (blue), ellipsoid (green), small circular (yellow), large circular (purple) FAs in the middle and FRET efficiency images in the bottom row. **(B)** FA count per FOV. **(C)** FA alignment within a given FOV (non-circular FAs only). **(D)** Stacked histogram representing the fraction of each morphological group per condition. **(E)** Average vinculin intensity representing vinculin recruitment. **(F)** Average FRET efficiency indicative of tension load (N = 80, 80, 61, 60 for CT-1kPa, CT-10kPa, CCM2-1kPa, CCM2-10kPa respectively, across 5 or 4 biological replicates). Each data point represents an average value per FOV which includes multiple cells. **(G-L)** Analysis of AJs using the VE-cadherin tension sensor (VECadTS) **(G)** Representative images of HBMVEC monolayers stably expressing VECadTS on 1 and 10 kPa collagen-coated PAA gels, grouped by control (siCT) and siCCM2-depleted samples. Donor fluorescence intensity images on the top row, morphology classification of mesh-like (blue) and linear (green) AJs in the middle and FRET efficiency images on the bottom row. **(H)** Average cell elongation calculated through the major and minor axis ratio of every cell. **(I)** Average AJ thickness. **(J)** Stacked histogram representing the fraction of each morphological group per condition. **(K)** Average VE-cadherin intensity representing VE-cadherin recruitment. **(L)** Average FRET efficiency indicative of tension load (N = 60, 60, 76, 79 for CT-1kPa, CT-10kPa, CCM2-1kPa, CCM2-10kPa respectively, across 4 or 5 biological replicates). Each data point represents an average value per FOV which includes multiple cells. All violin scatterplots are superplots distinguishing between technical replicates of each biological replicate. Statistical analysis between all groups was performed using linear mixed-effects model on the whole dataset (see Methods). *p < 0.05, **p < 0.01, ***p < 0.001.

Consistent with the well-established sensitivity of endothelial cells to ECM stiffness, both control and CCM2-depleted cells formed more FAs on 10 kPa than on 1 kPa substrates (Figure 4B). FA alignment was not significantly affected by substrate stiffness in either condition (Figure 4C). However, stiffness altered the distribution of FA morphologies in control cells, with fewer linear adhesions and more small circular adhesions on 10 kPa substrates. This redistribution was not observed in CCM2-depleted cells (Figure 4D; Figure S20A).

Increasing substrate stiffness also promoted vinculin recruitment to FAs in both control and CCM2-depleted cells (Figure 4E). At the molecular level, this was accompanied by an increase in vinculin tension on the stiffer substrates in both conditions (Figure 4F). Consistent with these observations, F-actin staining revealed higher actin intensity on 10 kPa than on 1 kPa substrates (Figure S21), indicating that increased vinculin recruitment and molecular tension occurred alongside increased F-actin content. The effects of substrate stiffness on vinculin recruitment and tension were detected across all FA categories (Figure S20B-C). Of note, differences between adhesion categories were less pronounced on 1 kPa than on 10 kPa substrates (Figure S22).

CCM2 depletion modified several aspects of the FA response to substrate stiffness. The increase in FA number from 1 to 10 kPa tended to be larger in CCM2-depleted than in control cells (Figure 4B) and FAs were significantly more aligned in CCM2-depleted cells on 10 kPa substrates compared to control cells (Figure 4C). In contrast, the distribution of FA morphologies remained largely unchanged with substrate stiffness in CCM2-depleted cells, unlike the redistribution observed in control cells (Figure 4D). Vinculin density and tension showed comparable stiffness-dependent increases in control and CCM2-depleted cells (Figure 4E,F). Vinculin recruitment nevertheless remained significantly higher in CCM2-depleted cells at both stiffnesses (Figure 4E).

Collectively, these results demonstrate that ECM stiffening promotes FA assembly, vinculin recruitment, and increased molecular tension in both control and CCM2-depleted endothelial cells. However, CCM2 loss modifies several aspects of this response. The increase in FA number is more pronounced, differences in FA alignment emerge at higher stiffness, and the stiffness-dependent redistribution of FA morphologies observed in control cells is lost. CCM2-depleted cells also show increased vinculin recruitment compared with controls, while vinculin tension remains comparable between the two groups. Thus, CCM2-deficient cells retain their ability to reinforce cell-ECM adhesions in response to matrix stiffness, but differ from control cells in how this response is reflected in FA organization.

In contrast to FAs, AJ organization was only modestly affected by substrate stiffness (Figure 4G). Cell shape (Figure 4H), cell size (Figure S23), AJ thickness (Figure 4I), and the distribution of linear and mesh-like junctional regions (Figure 4J; Figure S24A) did not change significantly between 1 and 10 kPa substrates in either control or CCM2-depleted cells. CCM2-depleted cells remained more elongated and larger than control cells at both stiffnesses (Figure 4H; Figure S23). Of note, differences in junction thickness and morphology between control and CCM2-depleted cells observed on 1 kPa substrates were no longer apparent on 10 kPa substrates (Figure 4I-J). This suggests that CCM2 depletion may limit the remodeling of junction thickness and morphology in response to changes in matrix stiffness.

Despite the limited effect of stiffness on AJ organization, VE-cadherin density increased from 1 kPa to 10 kPa in both control and CCM2-depleted cells (Figure 4K; Figure S24B). This effect tended to be larger in control cells, suggesting a reduced stiffness-dependent response in VE-cadherin recruitment following CCM2 depletion (Figure 4K). VE-cadherin tension also increased significantly with stiffness in control cells, whereas no significant change was detected in CCM2-depleted cells, especially for linear regions (Figure 4L; Figure S24C). Analysis of the individual junctional morphologies showed comparable VE-cadherin density between linear and mesh-like regions, while linear regions displayed lower tension in CCM2-depleted cells (Figure S25).

Collectively, these results indicate that AJ architecture is relatively insensitive to substrate stiffness, whereas VE-cadherin recruitment and molecular tension remain stiffness-responsive. This response is more evident in control endothelial cells, with CCM2 depletion attenuating the stiffness-dependent VE-cadherin recruitment and preventing a significant increase in VE-cadherin tension.

Together, these findings demonstrate that substrate stiffness influences endothelial adhesion mechanics, and that this response is altered by CCM2 loss. In control endothelial monolayers, increased substrate stiffness promoted coordinated increases in vinculin and VE-cadherin recruitment and molecular tension, reinforcing both cell-ECM and cell-cell adhesions. CCM2-depleted cells retained the stiffness-dependent increases in vinculin recruitment and tension at FAs, whereas increasing stiffness produced a smaller increase in VE-cadherin recruitment and no significant increase in VE-cadherin tension at AJs. Thus, increasing matrix stiffness further favors cell-ECM over cell-cell force transmission in CCM2-deficient endothelial cells, enhancing the mechanical imbalance associated with CCM2 loss.

## Discussion

### Disrupted endothelial force balance following CCM2 loss

Previous work using traction force microscopy (TFM) established that CCM-deficient endothelial cells exhibit altered force transmission at both cell-ECM and cell-cell interfaces. Isolated or paired CCM-deficient endothelial cells exert increased traction on the ECM (Faurobert et al., 2013; Shapeti et al., 2024) and higher intercellular tension at cell-cell contacts (Lisowska et al., 2018). However, these forces were examined separately and at the cellular or substrate level, leaving unresolved how CCM loss affects the balance of force transmission between the two adhesion systems within an endothelial monolayer. Here, we addressed this question by combining molecular tension measurements with quantitative analysis of adhesion organization in steady-state confluent monolayers. Using vinculin and VE-cadherin tension sensors, we could directly compare molecular loading at focal adhesions (FAs) and adherens junctions (AJs) and determine how CCM2 loss redistributes force between cell-ECM and cell-cell adhesions.

The ability to compare molecular tension across both adhesion systems extends previous applications of FRET-based tension sensors. Since its original development (Grashoff et al., 2010), the vinculin tension sensor has primarily been applied in single and/or migrating cells to study FA dynamics, rather than in confluent, steady-state monolayers relevant to tissue-level mechanics (Aytekin, Kimps, et al., 2026; Canever et al., 2026; Hernandez-Varas et al., 2015; LaCroix et al., 2018; Rothenberg et al., 2018; Sarangi et al., 2017; Shoyer et al., 2023). Applications of the VE-cadherin tension sensor have been more limited, including measurements of endothelial responses to shear stress (Conway et al., 2013, 2017), vascular development in zebrafish (Lagendijk et al., 2017) and leukocyte diapedesis (Arif et al., 2021; Uttekar et al., 2026). Combining VE-cadherin and vinculin tension sensors within the same steady-state, disease-relevant endothelial monolayer provides a direct comparison of cell-cell and cell-ECM molecular tension under identical experimental conditions, which has rarely been achieved within a single system.

This comparison revealed that CCM2 depletion redistributes molecular tension away from VE-cadherin at cell-cell junctions and toward vinculin at cell-ECM adhesions. CCM2-depleted cells formed more FAs, recruited more vinculin to these adhesions and exhibited higher molecular tension across vinculin. This distribution of increased molecular loading over more FAs and more vinculin molecules provides a potential molecular basis for the increased cell-ECM traction previously observed with TFM (Faurobert et al., 2013; Shapeti et al., 2024). At the cell-cell interface, however, CCM2 depletion produced thinner junctions, a shift from mesh-like toward linear morphology and reduced tension across VE-cadherin. The lower VE-cadherin tension contrasts with the higher intercellular tension previously measured using cell-cell TFM of isolated cell pairs (Lisowska et al., 2018), suggesting that this readout does not fully capture the molecular load transmitted specifically through VE-cadherin at adherens junctions.

These opposing responses at the two adhesion systems suggest that the mechanical consequences of CCM2 loss are better understood as a disruption of a coupled force-transmission system than as independent cell-cell and cell-ECM phenotypes. In previous studies, CCM proteins were found to engage with both cell-cell and cell-ECM adhesion machinery, and their loss has been associated with destabilized cell-cell junctions and reinforced cell-matrix engagement (Faurobert et al., 2013; Glading et al., 2007; Lisowska et al., 2018; Stockton et al., 2010; Whitehead et al., 2009). Mechanical coupling between cell-cell and cell-ECM adhesions is also well established as essential for tissue homeostasis (Andresen Eguiluz et al., 2017; Krishnan et al., 2011; Maruthamuthu et al., 2011). Our data extend this concept to the molecular scale by showing that CCM2 loss shifts the balance of molecular force transmission from cell-cell toward cell-ECM adhesions. This suggests that the CCM complex does not just regulate the two adhesion systems independently, but actively preserves the force transmission balance between them (Figure 5), positioning the CCM complex as a central hub to coordinate endothelial force transmission.

**Figure 5.**
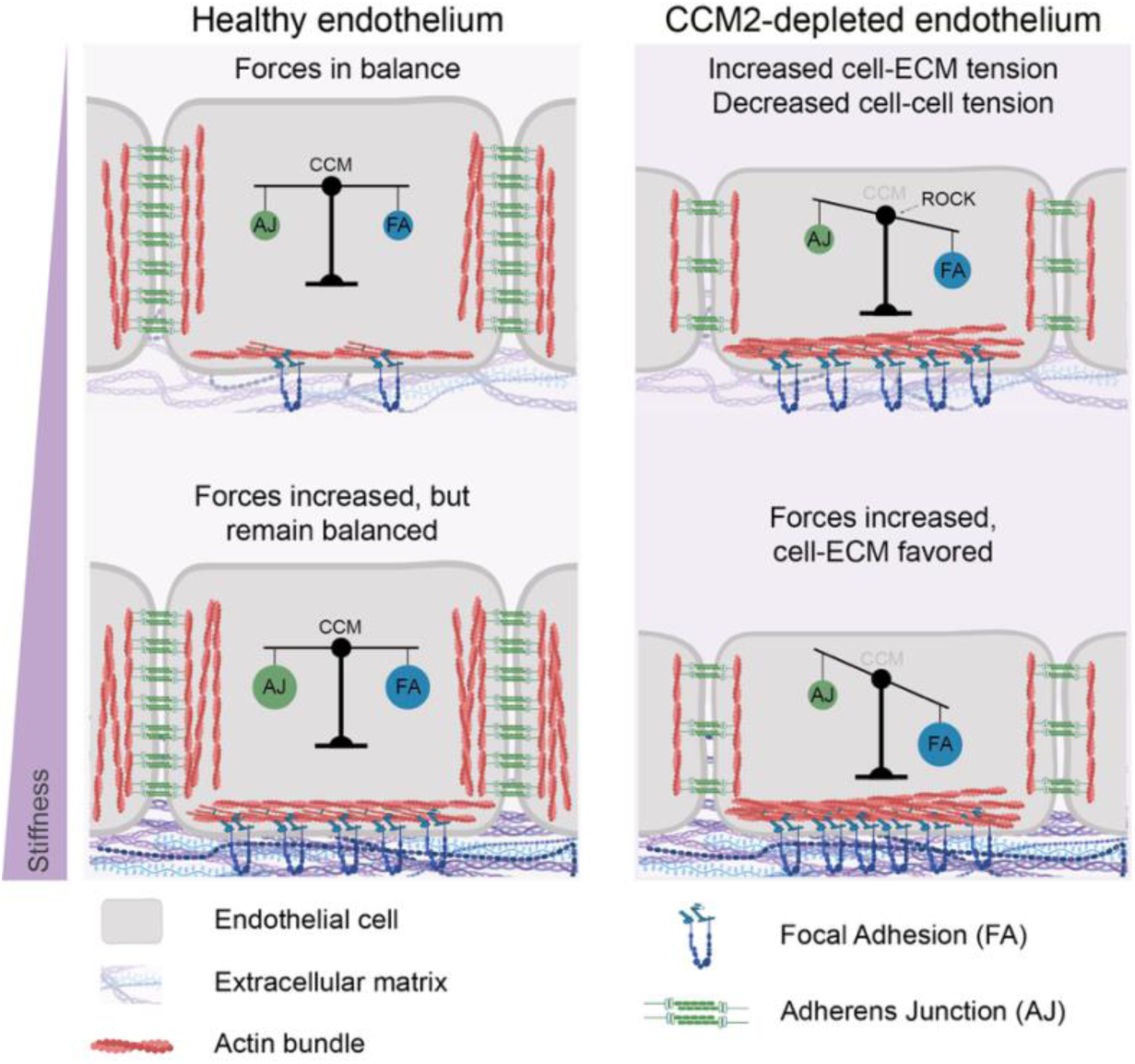
CCM2 loss redistributes, and ROCK sustains, the mechanical balance between cell-cell and cell-ECM adhesions on substrates with different stiffnesses. Summary model illustrating how CCM2 depletion, ROCK signaling, and substrate stiffness together shape the distribution of molecular tension between adherens junctions (AJ, green at cell-cell interface) and focal adhesions (FA, blue at cell-ECM interface) in the endothelial monolayer. **Top row:** In control (healthy) endothelium, tension across VE-cadherin and vinculin is balanced, reflecting coordinated force transmission between cell-cell and cell-ECM adhesions. CCM2 depletion shifts this balance: tension across vinculin at FAs increases while tension across VE-cadherin at AJs decreases, indicating a redistribution of mechanical load from cell-cell to cell-ECM adhesions rather than a uniform increase in contractility. Co-depletion of ROCK in CCM2-silenced cells on glass lowers tension at both adhesion types, with more limited effects on adhesion architecture, suggesting that ROCK activity is required to sustain, rather than to generate, this CCM2-associated tension imbalance. **Bottom row:** Increasing substrate stiffness raises tension at both adhesion systems in control monolayers while preserving their relative balance, whereas in CCM2-depleted cells stiffening preferentially reinforces FAs, amplifying the shift of mechanical load toward the cell-ECM interface. This schematic represents a proposed working model of how CCM2 loss, ROCK signaling, and matrix stiffness jointly shape mechanical crosstalk between cell-cell and cell-ECM adhesions in the endothelium.

At the cell-ECM interface, the increased FA formation, vinculin recruitment and molecular tension observed following CCM2 depletion are consistent with known effects of CCM loss on integrin activation and cytoskeletal organization. CCM2 loss destabilizes ICAP-1, relieving its inhibition of β1-integrin, and promoting both integrin-dependent FA assembly and cellular tractions (Faurobert et al., 2013; Millon-Frémillon et al., 2008). This increased cell-matrix engagement is also reflected in the pronounced cell elongation and FA alignment observed here, consistent with the integrin-driven, shear stress-like phenotype previously described in CCM2-deficient endothelial cells (Jilkova et al., 2014). More recently, the CCM complex has also been linked to FA and stress fiber organization through talin rod domain-containing protein 1 (TLNRD1), whose silencing increases cell spreading, stress fibers and paxillin-positive adhesion coverage (Ball et al., 2024). The distinct tension levels observed across FA morphologies provide an additional dimension to this mechanically activated state. In particular, CCM2 depletion increased the proportion of large circular FAs, which showed the highest vinculin loading, consistent with the complex, size-dependent tension-area FA relationship (Beningo et al., 2001; Hernandez-Varas et al., 2015; Tan et al., 2003). This can be better understood in the context of the ROCK-dependent senescence-associated secretory phenotype (SASP) reported in CCM-deficient endothelial cells, characterized by increased cell spreading, stress-fiber formation, and tractions, all of which promote persistent interactions with and remodeling of the surrounding ECM (Brauer et al., 2022; Chala et al., 2021; Vannier et al., 2021).

ROCK activity contributes to both sides of this imbalance (Figure 5). At FAs, co-depletion of CCM2 with either ROCK isoform substantially reduced vinculin tension, while changes in FA architecture and vinculin recruitment were modest. This suggests that ROCK-dependent contractility regulates molecular loading more directly than adhesion assembly itself, consistent with its position downstream of the ICAP-1/β1-integrin axis. The limited effect on FA architecture may reflect additional, ROCK-independent regulation of the structural phenotype. One potential mechanism involves TLNRD, whose actin-bundling activity is inhibited by CCM2. Consistent with a ROCK-independent contribution, TLNRD1 silencing alters stress fibers and cell-ECM adhesions without changing myosin light chain (MLC) phosphorylation (Ball et al., 2024). At cell-cell contacts, ROCK depletion similarly reduced VE-cadherin tension, bringing FRET efficiency close to that of the unloaded control sensor, despite limited effects on junction architecture. This strong dependence of molecular tension on ROCK is consistent with the proposed role of CCM proteins as a biophysical signaling hub at endothelial junctions that locally constrains RhoA/ROCK activity (Rödel & Abdelilah-Seyfried, 2026). Active Rap1 recruits CCM1 to cell-cell junctions, where its association with β-catenin physically links the CCM complex to VE-cadherin (Glading et al., 2007). At the junctions, the CCM complex limits Rho/ROCK signaling and favors cortical, junction-stabilizing actin over transverse stress fibers (Glading et al., 2007; Pannekoek et al., 2014; Stockton et al., 2010; Whitehead et al., 2009; D. Yang et al., 2026; Yin et al., 2024). Loss of the CCM complex disrupts this cortical actin network, and may therefore reduce force transmission through VE-cadherin. Within this reduced load, mesh-like regions carried more tension than linear ones after CCM2 depletion (Figure 2N), unlike the reticular junctions they resemble, which are proposed to occupy low-actin, low-tension cell overlaps (Fernández-Martín et al., 2012).

In our system, ROCK1 and ROCK2 sustain molecular tension across both adhesion systems, while their effects on adhesion architecture are more isoform-dependent. Specifically, CCM2-ROCK1 co-depletion had stronger effects on FA alignment, FA morphology and cell elongation than CCM2-ROCK2 co-depletion, in line with proposed complementary roles for the two isoforms in cytoskeletal remodeling and junctional mechanics (Beckers et al., 2015; Lisowska et al., 2018). The morphological rescue we observe following ROCK1 depletion was more modest than previously reported by Lisowska et al. (2018) (Lisowska et al., 2018). This difference might reflect variations in cell type, culture conditions or duration of depletion, as well as the co-expression of the tension sensor and endogenous protein in our system. Despite these isoform-specific effects on architecture, both ROCK1 and ROCK2 contribute similarly to maintaining molecular tension across both cell-cell and cell-ECM adhesions.

### Matrix stiffness amplifies the mechanical imbalance induced by CCM2 loss

The force balance between cell-cell and cell-ECM adhesions is also shaped by the mechanical properties of the ECM. This is particularly relevant in CCM, where growing evidence links ECM remodeling to disease progression (Faurobert et al., 2013; Hong et al., 2020; Shapeti et al., 2024; Vannier et al., 2021; X. Yang et al., 2026; Zhou et al., 2015). We therefore studied how substrate stiffness influences the force balance disrupted by CCM2 loss, using stiffness values estimated for healthy brain and CCM lesion tissue. In control monolayers, increasing stiffness enhanced both vinculin and VE-cadherin recruitment and molecular tension. This coordinated reinforcement of cell-ECM and cell-cell adhesion is consistent with a mechanical balance in which increased cell-matrix loading is matched by reinforcement of the cell-cell interface, preserving monolayer integrity (Andresen Eguiluz et al., 2017; Krishnan et al., 2011; Maruthamuthu et al., 2011).

CCM2-depleted cells retained the ability to sense matrix stiffness, but this response was no longer coordinated across the two adhesion systems. Increasing stiffness reinforced vinculin recruitment and tension at FAs, whereas the response at AJs was weaker, with a smaller increase in VE-cadherin recruitment and no significant increase in VE-cadherin tension. Matrix stiffening therefore does not simply amplify the effects of CCM2 loss at both interfaces. Instead, it enhances the asymmetry between the two adhesion systems, further shifting force transmission toward the cell-ECM interface, increasing the mechanical imbalance between the two adhesion systems (Figure 5). These results support a role for the CCM complex not only in regulating force transmission at individual adhesions, but also in coordinating how endothelial cells adjust the balance between cell-cell and cell-ECM forces in response to their mechanical environment.

Increasing matrix resistance is known to promote force generation and stabilization of integrin-based adhesions (Elosegui-Artola et al., 2016; Kim & Wirtz, 2013; Pelham & Wang, 1997). However, CCM2-depleted cells showed enhanced vinculin recruitment even on soft 1 kPa substrates, reaching levels comparable to those of control cells on stiffer 10 kPa substrates. Combined with the absence of a stiffness-dependent change in FA shape distribution, this suggests that CCM2-depleted cells may respond to matrix at a lower mechanical threshold. This mechanically primed state at the cell-ECM interface may be another consequence of the β1-integrin activation following disruption of the CCM2/ICAP-1 axis, with effects that could extend beyond the cell-ECM interface itself (Faurobert et al., 2013; Millon-Frémillon et al., 2008).

This primed state raises the possibility of a self-reinforcing mechanical loop in which CCM2-associated β1-integrin hyperactivation and increased cell-ECM force promote matrix remodeling and stiffening, which in turn reinforces cell-ECM adhesion, leading to a positive feedback loop. This possibility is supported by histopathological reports of fibrosis, collagen deposition, hyalinization, and calcification in CCM lesions (Frischer et al., 2008; Ren et al., 2025), features commonly associated with tissue stiffening. Such a mechanism may be particularly relevant to the mosaic nature of CCM lesions, in which mutant cells coexist with and recruit neighboring wild-type endothelial cells (Detter et al., 2018; Malinverno et al., 2019; Rath et al., 2020; Shapeti et al., 2024). Local matrix remodeling driven by CCM-deficient cells could therefore alter the mechanical environment experienced by neighboring wild-type cells, potentially promoting FA reinforcement as previously observed *in vitro* (Faurobert et al., 2013).

## Conclusion

By combining molecular tension sensing with quantitative analysis of adhesion organization, we established an integrated approach to investigate how molecular forces, cellular architecture, and the mechanical microenvironment interact in disease. Applied to CCM, this approach revealed that the CCM complex plays a dual mechanical role in the endothelium by balancing force transmission between cell-cell and cell-ECM adhesions and coordinating how this balance is adjusted in response to changes in the mechanical microenvironment. Matrix stiffness emerges as a potential amplifier of the force imbalance induced by CCM2 loss, suggesting that altered mechanotransduction and the extracellular environment may reinforce each other during disease progression. The resulting shift toward cell-ECM force transmission, together with the reduced coordination of cell-cell and cell-ECM responses to matrix stiffness, may compromise endothelial cohesion and adaptation to the local mechanical environment, contributing to vascular instability in CCM.

Future studies using dynamically tunable matrices and time-resolved tension measurements could test whether altered cell-ECM force transmission drives matrix remodeling and, in turn, whether these matrix changes further reinforce the mechanical phenotype of CCM-deficient cells. Incorporating physiological flow and mosaic endothelial models could further reveal how this force imbalance develops and propagates within the more complex mechanical environment of CCM lesions.

## Material & methods

### Molecular cloning

The parental plasmids used for molecular cloning were obtained from Addgene. pcDNA3.1-VinTS (Addgene #111764) was a gift from Brenton Hoffman (LaCroix et al., 2018) and pLPCX-VECadTS (Addgene #45848) was a gift from Martin Schwartz (Conway et al., 2013). The constructs containing FL (Addgene #101170) and HP35st (Addgene #101251) elastic linkers were a gift from Carsten Grashoff (Austen et al., 2015; Ringer et al., 2017). The Clover construct (Addgene #40259) was a gift from Michael Lin (Lam et al., 2012). The pHR backbone (Addgene #79129) was a gift from Wendell Lim (Morsut et al., 2016).

Viral vector constructs were generated through a series of cloning steps. First, the tension sensor modules (TSMods; Figure 1A) containing Clover as donor, an elastic linker (F40, FL or HP35st) and mScarlet-I as acceptor were generated through PCR amplification (Phusion^TM^ Thermo Scientific) using primers containing the required overhangs (list in Supplementary note 1). To reduce repetitive nucleotide sequence content, the F40 elastic linker was synthesized as a human codon optimized gBlock (IDT; Supplementary note 2). The resulting PCR products were assembled through Gibson assembly (GeneArt™ Gibson Assembly HiFi Master Mix Thermo Scientific), into a pcDNA3.1 backbone enzymatically digested with NheI and EcoRI (FastDigest Thermo Scientific).

Before inserting the new TSMods into the full tension sensors, the VE-cadherin sensor was adapted to match the cloning design of the pcDNA3.1-VinTS construct (*NheI* – head – Clover *(with KpnI)* – linker – mScarlet-I *(with Bsp1407I)* – tail – *EcoRI*). This was accomplished in two steps. First, pcDNA3.1-VECadTL was generated by inserting the VE-cadherin head amplified from pLPCX-VECadTS together with an amplified TSMod into NheI/EcoRI-digested pcDNA3.1 by Gibson assembly. Second, the VE-cadherin tail was amplified from pLPCX-VECadTS in two fragments and inserted by Gibson assembly into Bsp1407I/EcoRI-digested pcDNA3.1-VECadTL. This strategy removed an additional Bsp1407I restriction site present in the original VE-cadherin tail and corresponded to our common cloning design for the vinculin and VE-cadherin sensors (primer list in Supplementary note 1).

The F40/FL/HP35st TSMods were then inserted into the vinculin (Vin) and VE-cadherin (VECad) constructs using KpnI/Bsp1407I or KpnI/EcoRI digestion and ligation. This generated the corresponding tension sensor (VinTS and VECadTS) and tailless control (VinTL and VECadTL) constructs respectively containing each of the three elastic linkers (VinTS-F40, VinTS-FL, VinTS-HP35st; VECadTS-F40, VECadTS-FL, VECadTS-HP35st; VinTL-F40, VinTL-FL, VinTL-HP35st; VECadTL-F40, VECadTL-FL, VECadTL-HP35st).

Finally, the pHR backbone was modified to facilitate transfer of the tension sensors and to provide an empty vector control for background FLIM measurements. The original backbone was digested with NheI and EcoRI and assembled through Gibson with a synthesized SFFV-EcoRI-NheI-WPRE-3’LTR cassette (Genscript; Supplementary note 2) with the original NheI and EcoRI sites destroyed during assembly. All tension sensors were then transferred into this modified pHR backbone by NheI/EcoRI digestion and ligation. The resulting viral constructs were pHR_VinTS, pHR_VinTL, pHR_VECadTS, pHR_VECadTL, each containing an F40, FL or HP35st elastic linker between the minimal Clover and mScarlet-I fluorophore.

### Cell culture and viral transduction

Pooled Human Brain Microvascular Endothelial Cells (HBMVECs) were obtained from PeloBiotech (REF #PB-CH-110-4011) and grown in EGM-2 MV microvascular complete medium (Lonza, REF #CC-3156 and #CC-4147) supplemented with 100 U/mL penicillin and 100 μg/mL streptomycin on collagen type I-coated dishes (100 µg/mL, 30 min incubation at 37 °C) (Thermo Fisher Scientific, REF #A1048301). Human Embryonic Kidney 293T cells (HEK-293T) were obtained from ATCC (REF #CRL-3216) and maintained in Dulbecco’s Modified Eagle Medium (DMEM) with 4.5 g/L D-glucose (Gibco, REF #31053028) supplemented with 10% v/v fetal bovine serum (Gibco, REF #F7524), 1% GlutaMax (Gibco, REF #35050038) and 50 µg/mL gentamicin (Carl Roth, REF #2475.1). Cells were grown at 37 °C with 5% CO2. The use of these commercially sourced primary human cells was approved by the Ethics Committee Research UZ/KU Leuven (study number S71563).

24 h after seeding 1.1 x 10^6 cells per well in a 6-well plate (Sarstedt, REF #83.3920), HEK-293T cells were transfected using Turbofect™ Transfection Reagent (Thermo Scientific, REF #R0533) according to the manufacturer’s instructions. Per well, 950 ng of pCMVR8.74 packaging (gift from Didier Trono, Addgene #22036), 570 ng of pMD2.G envelope (gift from Didier Trono, Addgene #12259) and 475 ng of a lentiviral pHR plasmid carrying one of the tension sensor plasmids previously mixed with 8 µL Turbofect were added to 400 µL Opti-MEM (Opti-MEM™ Reduced Serum Medium Gibco, REF #31985062). The mixture was incubated for 15 minutes at room temperature and then added dropwise to the cells. 6 h after transfection, the medium was replaced with fresh DMEM. After 24 h, 3.5 x 10^4 HBMVECs were seeded on collagen-coated 6-well plates. Lentivirus production proceeded for 48 h in the HEK-293T, after which the virus-containing supernatant was harvested and filtered through a 0.45 µm syringe filter (SFCA Syringe Filters Corning™, REF #10002471 and Luer lock syringes, Fisher Scientific REF #15879152). After addition of 10 µg/mL polybrene (Sigma-Aldrich, REF #TR-1003), half the collected virus volume per well was added dropwise to the HBMVEC cells and incubated for 8 h, after which the cells were washed three times with 1x PBS (Gibco, REF #14200075). Over the following days, three additional washes were performed before the cells were frozen in freezing medium (5% DMSO (Fisher Bioreagents, REF #BP231-100) and 95% FBS) or reseeded for subsequent treatments.

### Transient transfection with siRNA

For 6-well plates (Cellvis, REF #P06-14-0-N), 2 x 10^5 HBMVECs were seeded 24 h before the first transfection. Next, the cells were transfected twice at 24 h intervals with 0.72 µL (100 µM) of siRNA in 300 µL opti-MEM and 6 µL Lipofectamine RNAiMAX (Invitrogen^TM^, REF #13778150) in 300 µL opti-MEM. The lipofectamine was added to the siRNA and incubated for 15-30 minutes before being added dropwise to the cells. The medium was replaced after 4 h incubation. For double transfections, 0.72 µL (100 µM) of each siRNA was used with the same volume of Lipofectamine RNAiMAX. The siRNA duplexes used: control (siCT) (Dharmacon ON-TARGETplus Non-targeting Control siRNAs, REF #D-001810-01), siCCM2 (Dharmacon ON-TARGETplus SMARTpool, REF #L-014728-01), siROCK1 (Dharmacon ON-TARGETplus SMARTpool, REF #L-003536-00) and siROCK2 (Dharmacon ON-TARGETplus SMARTpool, REF #L-004610-00). Experiments can be downscaled to 24-well (Cellvis, REF #P24-0-N) plates for ∼25% the volumes.

### Substrate preparation (PAA) and cell transfer

PAA hydrogels of different stiffnesses were prepared based on previously published protocols (Cresens et al., 2026; Aytekin et al., 2026). Glass-bottom dishes (Cellvis, REF #D35-20-0-N) were pre-treated with bind-silane (Serva, REF # 28739.01) and equipped with a 9 mm diameter 120 µm thick spacer (Secure-Seal™ Grace Bio-Labs, REF #654002). Cover glasses of 12 mm diameter (Fisher Scientific, REF #NC9216450) were also pre-treated with Sigma-Coat (Sigma-Aldrich, REF # SL2-25ML). Next, gels with a Young’s modulus of 1 kPa or 10 kPa were prepared by mixing 40% acrylamide (AAm) (Sigma-Aldrich, REF #A4058) and 2% bisacrylamide (MBAA) (Sigma-Aldrich, REF #M1533) in dH2O (for exact volumes see Table 1) on ice. The polymerization of the gel was initiated by ammonium persulfate (10% APS) (Bio-Rad, REF #1610700EDU) and catalyzed by N,N,N’,N’-tetramethylethylenediamine (TEMED) (Sigma-Aldrich, REF #T7024). The mixture was immediately transferred to the previously treated glass-bottom dishes and flattened with the siliconized cover glass and left to polymerize at 37 °C for 10 minutes. Polymerized gels were washed three times with 1x PBS. Afterwards, the gel surface was functionalized by two rounds of 0.2 mg/mL Sulfo-SANPAH (Sigma-Aldrich, REF #803332) treatment under 365 nm UV irradiation for 15 minutes. After three washes with 1x PBS, gels (and corresponding glass samples) were incubated for 30 minutes at 37 °C with 25 µg/mL collagen type I diluted in 0.1% v/v acetic acid (Acros Organics, REF #64-19-7) at pH 3.4 for homogeneous coating. This was followed by a last washing round with 1x PBS. Cells were transferred to the PAA gels 24 h after the last siRNA transfection round by reseeding 2 x 10^5 treated HBMVECs on the gel samples.

**Table 1:**
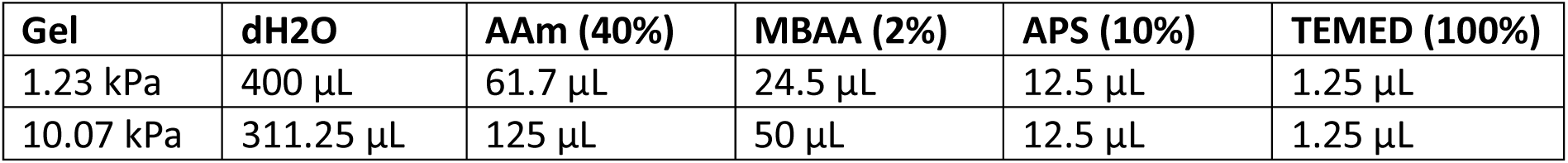
PAA composition adapted from (Cresens et al., 2026).

### Sample fixation and actin staining

72 h after the last siRNA transfection round, all samples were fixed with 4% paraformaldehyde (PFA) (Thermo Fisher Scientific, REF #28908) for 15 minutes at 37 °C. Cells were washed three times with 1x PBS and stored at 4°C. When actin staining was necessary, cells were incubated with 100 nM of SiR-actin (SpiroChrome, REF #SC001) at room temperature for 15 minutes followed by a single washing step with 1x PBS right before imaging.

### Quantitative RT-PCR

RNA was isolated from cells with the NucleoSpin RNA II kit (Macherey-Nagel, REF #740955) following the manufacturer’s instructions. cDNA was synthesized from 500 ng of RNA using the High-Capacity cDNA Reverse Transcription Kit (ThermoFisher, REF # 4368814). Quantitative real-time PCR was run with iTaq Universal SYBR Green Supermix (Bio-Rad, REF #L001752B) on a C-1000 Touch thermal cycler (Bio-Rad Laboratories) and melt curves were analyzed in CFX Manager software (Bio-Rad Laboratories). We used two housekeeping genes (GAPDH and RANBP1) to evaluate the expression levels of CCM2, ROCK1 and ROCK2 in the siRNA-silenced HBMVEC samples. Primer sequences for all PCR reactions are provided in Supplementary note 3.

### Fluorescence microscopy imaging

FLIM-FRET data were collected on a Leica TCS SP8 confocal microscope fitted with the FALCON (FAst Lifetime CONtrast) module. All measurements were acquired at room temperature on fixed samples imaged in PBS. A 63× HC PL APO water-immersion objective (NA 1.20) was used, and images of 512 × 512 pixels were recorded with an image size of 123.26 x 123.26 µm (pixel size 240.7 nm). Unidirectional scanning was set to 50 Hz, with a pixel dwell time of 30 µs. Laser power of 1.35 µW (488 nm) was measured using a Thorlabs PM100USB power meter. Clover was excited at 488 nm and its emission was recorded between 495–540 nm with a HyD SMD detector in counting mode. mScarlet-I was excited at 561 nm, and emission between 570–635 nm was collected with a HyD SMD detector in counting mode. Each FLIM acquisition consisted of six donor frames and one acceptor frame, using a pulsed laser with 20 MHz repetition rate. For each condition 60 to 217 FOVs were obtained. Clover-only and samples transduced with empty pHR vector were measured to obtain the donor-only and background values.

Excitation & emission SiR-actin!

### FLIM-FRET analysis

For the FRET analysis, the fluorescence decay of every pixel was converted to phasor space with the integrated Phasor module of the SP8 FALCON. Briefly, the full decay *I(t)* was Fourier transformed into a real component (*G*) and an imaginary component (*S*) according to:

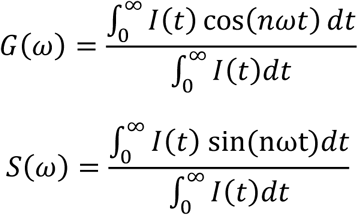

where n indicates the harmonic used (here 1), *ω* equals *2πf* and *f* is the repetition rate of the pulsed laser (here 20 MHz). The *G* and *S* values give the phasor coordinates of each pixel, which form a 2D histogram of the lifetime information contained in the image. The exported *G* and *S* coordinates were analyzed further with a custom MATLAB script for phasor-based FRET calculation (Aytekin, Kimps, et al., 2026; Coucke et al., 2023, 2024). The starting point of the FRET efficiency trajectory was set at the position on the phasor semicircle corresponding to the Clover fluorescence lifetime (∼3.0 ns), measured in donor-only HBMVECs expressing cytosolic Clover. The end point of the trajectory was set by the approximate coordinates of the background signal (*G = 0.5, S = 0.25*), as observed in HBMVECs transduced with the empty pHR vector. A 9 x 9 pixel median filter was applied to the *G* and *S* coordinate images before assigning FRET values, to reduce noise and smooth the resulting FRET efficiency distribution, as described similarly by Ranjit et al. (2018) (Ranjit et al., 2018). Each pixel of the image, represented by a point in phasor space, was then assigned the FRET value of the nearest point on the FRET trajectory, producing the FRET image.

### Quantitative image analysis of FA and AJ organization

The resulting intensity and FRET data were analyzed using an in-house custom MATLAB script (available on https://github.com/SarahVorsselmans). Segmentation of FAs and AJs was performed in Ilastik and further defined manually (e.g. to remove central VE-cadherin vesicles or out-of-focus structures on gels). The resulting segmentation was used to define a binary mask for each field of view (FOV). Background objects smaller than 0.4 µm^2^ and 1.16 µm^2^ in surface area were removed from the mask for vinculin and VE-cadherin, respectively. The acceptor-channel image was re-aligned to the donor channel and used for all subsequent intensity calculations. The non-segmented donor image was retained only for representative display.

For each FOV, FAs were identified as individual segmented objects and counted per FOV to obtain the FA count. Per-object shape descriptors were extracted by fitting an ellipse to each FA, finding its major and minor axis lengths. The ratio of minor to major axis (MinMajRatio) was used as a roundness index, ranging from ∼0 (linear) to 1 (fully round). FA alignment within a FOV was quantified by first excluding round objects (MinMajRatio ≥ 0.667), converting the orientation of the remaining elongated FAs to a −90° to 90° scale, calculating the mean FA angle per FOV, and then scoring how closely individual FA angles matched this mean to obtain a per-FOV alignment index.

FAs were further classified morphologically based on shape and size. Objects were first divided into elongated (MinMajRatio < 0.5) and round (MinMajRatio ≥ 0.5) structures. Elongated FAs were subclassified as linear (MinMajRatio < 0.3) or ellipsoid (0.3 ≤ MinMajRatio < 0.5). Round FAs were subclassified by area into small circular (< 1.44 µm^2^) and large circular (≥ 1.44 µm^2^) adhesions (Kanchanawong & Calderwood, 2022; Tan et al., 2003). For each morphological category, the fractional abundance, mean FRET efficiency, and mean intensity were calculated by isolating all pixels belonging to that category and calculating their average per FOV.

FRET efficiency and intensity were each also calculated at the whole-FOV level, as the mean of all valid (non-background) segmented pixels rather than per individual FA. Per-FOV averages were only calculated for FOVs containing at least 5 FAs. Since the FAs were rather small in comparison to e.g. fibroblasts, we did not investigate the FRET efficiency gradients over the FA and instead averaged them. Actin images were analyzed by first removing background pixels below a fixed intensity threshold of 20 (a.u.) followed by calculating the mean intensity of the remaining pixels per image.

Cell shape was quantified per cell by fitting an ellipse to each segmented cell area, from which major and minor axis lengths (converted to µm using the pixel size) and a minor:major axis ratio (roundness index) were calculated and averaged per FOV.

AJ thickness was measured along the cell-cell junction skeleton. Small segmented areas were removed and small gaps closed, after which the contour was smoothed. A padded Euclidean distance transform (EDT) was then applied, assigning each pixel the radius of the largest inscribed disk centered on it. A medial-axis skeleton was extracted from the smoothed mask and each skeleton pixel was assigned the corresponding EDT value. This was then doubled to convert radius to diameter and converted to µm. This yielded a thickness value for every skeleton pixel. The mean of these values gave a single per-FOV thickness estimate.

AJs were also classified morphologically similarly to the FA classification above. Junction masks were first split into individual segments by watershed sectioning of the distance transform, which separates morphologically distinct regions along a continuous junction while preserving small enclosed gaps as holes. Each segment was then scored for its local thickness (diameter, from the same EDT-based approach as above) and for the presence of enclosed holes, represented by the Euler number. Segments that were both thick (≥ 1 µm) and contained holes were classified as mesh-like junctions, while the remaining thinner and more uniform segments were classified as linear junctions. For each FOV, the fractional abundance of each category was calculated from the proportion of junction skeleton pixels assigned to it.

### Statistics and reproducibility

Statistical analyses were performed using linear mixed-effects models (LMMs) due to the unequal number of biological and technical replicates. This was implemented in MATLAB (fitlme) (available on https://github.com/SarahVorsselmans). For each quantified metric (FA count, FA alignment, cell elongation, junction thickness, fraction, average intensity, FRET efficiency), a single model was fit across the entire dataset simultaneously, containing all experimental conditions (control and CCM2-depleted cells, ROCK1/ROCK2 co-silencing, 1 and 10 kPa substrates, and tailless sensor controls), rather than fitting separate models per figure or per subset of conditions. The model was set to *metric ∼ condition + (1 | biological replicate)* where condition was a fixed factor encoding all experimental groups directly and biological replicate was included as a random intercept to account for the unbalanced number of observations across independent experiments. Models were fit by restricted maximum likelihood (REML) with effects coding of the fixed-effect design matrix. Prior to model fitting, technical replicates were averaged to a single value per condition per biological replicate to avoid overinflation. This mirrors the logic of "SuperPlots" while allowing formal mixed-model contrasts and correction for multiple comparisons. SuperPlots is a plotting approach that separates the spread of individual technical replicate measurements from the underlying biological replicate structure. In each SuperPlot every technical replicate is shown as a data point with its shape and color together identifying the biological replicate it came from. Each data point represents an average value per FOV which includes multiple cells. Averages per biological replicate are depicted as larger symbols with a black border. The violin plot lines represent the 75th percentile, median and 25th percentile.

Pairwise comparisons between conditions were performed as linear hypothesis tests on the fixed-effect coefficients of the fitted model (contrast testing), providing a p-value for each comparison. Certain pairwise comparisons were excluded from testing because they were not biologically or experimentally meaningful: (i) any comparison between the PAA gel and ROCK-depleted samples was not included; (ii) any comparison between a full-length tension sensor (TS) and its corresponding tailless (TL) control because these serve as an internal loading control rather than an experimental condition; and (iii) the "crossed" stiffness/genotype comparisons CT-1kPa-TS versus CCM2-10kPa-TS and CT-10kPa-TS versus CCM2-1kPa-TS, which would mix the effects of genotype and substrate stiffness and were therefore not useful as single-variable comparisons. Raw p-values were corrected for multiple comparisons using Benjamini-Hochberg false discovery rate (FDR). Significance was indicated as *p < 0.05, **p < 0.01, ***p < 0.001.

For comparisons of intensity and FRET efficiency between morphological categories (e.g. Figure 2G, 2N) within the same experimental condition, repeated-measures ANOVA (rmANOVA) was used on the averaged biological replicates instead of the LMM framework described above since these comparisons involved the same biological and technical replicates measured repeatedly across categories rather than between independent groups. Lastly, for the comparison of FRET efficiency between the full-length tension sensor (TS) and its corresponding tailless (TL) control for each elastic linker (Figure S1), a paired t-test was used instead.

## Code and data availability

All custom MATLAB code used for FLIM-FRET phasor calculation, image analysis and statistical testing is available at https://github.com/SarahVorsselmans [https://github.com/SarahVorsselmans/FRET-and-Morphology-at-FA-and-AJ.git]. The full dataset tables underlying the figures have been deposited on Zenodo [https://doi.org/10.5281/zenodo.23164604].

## Use of AI tools

Claude (Anthropic; Sonnet 3.7-5.5; accessed April-October 2026) was used to assist with writing MATLAB code for data analysis and with editing the manuscript text. Based on analysis systems specified by the authors, Claude generated and edited MATLAB scripts which were then reviewed, tested and validated by the authors who remain responsible for the final analysis pipeline. Claude was used to improve the language, flow and structure of text drafts written by the authors who also reviewed and revised all AI-assisted text, verified all statements and references and take full responsibility for the content of the manuscript. Claude was not used to generate, alter or interpret experimental data or images, nor did it contribute to the study design, results or conclusions.

## Acknowledgments and funding

The authors are grateful to Rik Nuyts for assistance with microscopy measurements and imaging optimization, and Hendrik Cuylaerts for assistance in construct generation. They are grateful to the following funding sources: S.R. and S.A. were supported by KU Leuven internal funding C14/22/085. S.R., S.V. and E.F. received funding from the Research Foundation Flanders (FWO) through a project with grant number G0C2422N. E.F. also acknowledges Ambition Internationale, Région AURA, France. In addition, S.A. is a recipient of FWO fellowships with grant number 1S95125N. S.V. acknowledges KU Leuven Facultaire Luik Onderzoeksfonds (FLOF) and EMBO Scientific Exchange Grant 10986 for financial support.

## Author contributions

S.V. designed, performed and interpreted experiments and data analysis. E.F. and S.R. conceptualized and supervised the study and designed and interpreted experiments. S.V. and S.A. generated the DNA constructs used in this study with input from C.O. The customized morphology algorithms were developed by S.V. with input from T.M.R. The manuscript was written by S.V. under supervision of S.R. and E.F. All authors contributed to the writing of the final version.

## References

Andresen Eguiluz, R. C., Kaylan, K. B., Underhill, G. H., & Leckband, D. E. (2017). Substrate stiffness and VE-cadherin mechano-transduction coordinate to regulate endothelial monolayer integrity. Biomaterials, 140, 45–57. 10.1016/J.BIOMATERIALS.2017.06.010

Arif, N., Zinnhardt, M., Nyamay’Antu, A., Teber, D., Brückner, R., Schaefer, K., Li, Y., Trappmann, B., Grashoff, C., & Vestweber, D. (2021). PECAM-1 supports leukocyte diapedesis by tension-dependent dephosphorylation of VE-cadherin. The EMBO Journal, 40(9), e106113. 10.15252/EMBJ.2020106113

Austen, K., Ringer, P., Mehlich, A., Chrostek-Grashoff, A., Kluger, C., Klingner, C., Sabass, B., Zent, R., Rief, M., & Grashoff, C. (2015). Extracellular rigidity sensing by talin isoform-specific mechanical linkages. Nature Cell Biology 2015 17:12, 17(12), 1597–1606. 10.1038/ncb3268

Ayata, C., Kim, H., Morrison, L., Liao, J. K., Gutierrez, J., Lopez-Toledano, M., Carrazana, E., Rabinowicz, A. L., & Awad, I. A. (2024). Role of Rho-Associated Kinase in the Pathophysiology of Cerebral Cavernous Malformations. Neurology: Genetics, 10(1), e200121. 10.1212/NXG.0000000000200121

Aytekin, S., Kimps, L., Coucke, Q., Linhares, D., Vorsselmans, S., Deodhar, S., Cardinaels, R., Cóndor, M., Barrasa-Fano, J., Van Oosterwyck, H., & Rocha, S. (2026). Linking molecular tension and cellular tractions: a multiscale approach to focal adhesion mechanics. Communications Biology, 9(1), 236. 10.1038/S42003-026-09514-0

Aytekin, S., Vorsselmans, S., Vankevelaer, G., Poedts, B., Hendrix, J., & Rocha, S. (2026). From Sensor Design to Force Maps: A Systematic Evaluation of FRET-based Vinculin Tension Sensors. BioRxiv, 2026.03.23.713753. 10.64898/2026.03.23.713753

Ball, N. J., Ghimire, S., Follain, G., Pajari, A. O., Wurzinger, D., Vaitkevičiūtė, M., Cowell, A. R., Berki, B., Ivaska, J., Paatero, I., Goult, B. T., & Jacquemet, G. (2024). TLNRD1 is a CCM complex component and regulates endothelial barrier integrity. Journal of Cell Biology, 223(9). 10.1083/JCB.202310030/276861

Beckers, C. M. L., Knezevic, N., Valent, E. T., Tauseef, M., Krishnan, R., Rajendran, K., Hardin, C. C., Aman, J., van Bezu, J., Sweetnam, P., van Hinsbergh, V. W. M., Mehta, D., & van Nieuw Amerongen, G. P. (2015). ROCK2 primes the endothelium for vascular hyperpermeability responses by raising baseline junctional tension. Vascular Pharmacology, 70, 45. 10.1016/J.VPH.2015.03.017

Beningo, K. A., Dembo, M., Kaverina, I., Small, J. V., & Wang, Y. L. (2001). Nascent Focal Adhesions Are Responsible for the Generation of Strong Propulsive Forces in Migrating Fibroblasts. The Journal of Cell Biology, 153(4), 881. 10.1083/JCB.153.4.881

Bergametti, F., Denier, C., Labauge, P., Arnoult, M., Boetto, S., Clanet, M., Coubes, P., Echenne, B., Ibrahim, R., Irthum, B., Jacquet, G., Lonjon, M., Moreau, J. J., Neau, J. P., Parker, F., Tremoulet, M., & Tournier-Lasserve, E. (2005). Mutations within the programmed cell death 10 gene cause cerebral cavernous malformations. American Journal of Human Genetics, 76(1), 42–51. 10.1086/426952

Borikova, A. L., Dibble, C. F., Sciaky, N., Welch, C. M., Abell, A. N., Bencharit, S., & Johnson, G. L. (2010). Rho kinase inhibition rescues the endothelial cell cerebral cavernous malformation phenotype. Journal of Biological Chemistry, 285(16), 11760–11764. 10.1074/jbc.C109.097220

Brandon, K. D., Frank, W. E., & Stroka, K. M. (2024). Junctions at the crossroads: the impact of mechanical cues on endothelial cell-cell junction conformations and vascular permeability. Https://Doi.Org/10.1152/Ajpcell.00605.2023, 327(4), C1073–C1086. 10.1152/AJPCELL.00605.2023

Brauer, E., Lange, T., Keller, D., Görlitz, S., Cho, S., Keye, J., Gossen, M., Petersen, A., & Kornak, U. (2022). Dissecting the influence of cellular senescence on cell mechanics and extracellular matrix formation in vitro. Aging Cell, 22(3), e13744. 10.1111/ACEL.13744

Budday, S., Nay, R., de Rooij, R., Steinmann, P., Wyrobek, T., Ovaert, T. C., & Kuhl, E. (2015). Mechanical properties of gray and white matter brain tissue by indentation. Journal of the Mechanical Behavior of Biomedical Materials, 46, 318–330. 10.1016/J.JMBBM.2015.02.024

Canever, H., Lachuer, H., Delaunay, Q., Sipieter, F., Audugé, N., Girard, P. P., & Borghi, N. (2026). Collective directional memory controls the range of epithelial cell migration. ELife, 15. 10.7554/ELIFE.110739.1

Chala, N., Moimas, S., Giampietro, C., Zhang, X., Zambelli, T., Exarchos, V., Nazari-Shafti, T. Z., Poulikakos, D., & Ferrari, A. (2021). Mechanical Fingerprint of Senescence in Endothelial Cells. Nano Letters, 21(12), 4911–4920. 10.1021/ACS.NANOLETT.1C00064

Clatterbuck, R. E., Eberhart, C. G., Crain, B. J., & Rigamonti, D. (2001). Ultrastructural and immunocytochemical evidence that an incompetent blood-brain barrier is related to the pathophysiology of cavernous malformations. Journal of Neurology, Neurosurgery, and Psychiatry, 71(2), 188. 10.1136/JNNP.71.2.188

Conway, D. E., Breckenridge, M. T., Hinde, E., Gratton, E., Chen, C. S., & Schwartz, M. A. (2013). Fluid shear stress on endothelial cells modulates mechanical tension across VE-cadherin and PECAM-1. Current Biology : CB, 23(11), 1024. 10.1016/J.CUB.2013.04.049

Conway, D. E., Coon, B. G., Budatha, M., Arsenovic, P. T., Orsenigo, F., Wessel, F., Zhang, J., Zhuang, Z., Dejana, E., Vestweber, D., & Schwartz, M. A. (2017). VE-cadherin phosphorylation regulates endothelial fluid shear stress responses through the polarity protein LGN. Current Biology : CB, 27(14), 2219. 10.1016/J.CUB.2017.06.020

Coucke, Q., Parveen, N., Fernández, G. S., Qian, C., Hofkens, J., Debyser, Z., & Hendrix, J. (2023). Particle-based phasor-FLIM-FRET resolves protein-protein interactions inside single viral particles. Biophysical Reports, 3(3), 100122. 10.1016/j.bpr.2023.100122

Coucke, Q., Vorsselmans, S., Aytekin, S., Huysecom, S., Hofkens, J., Hendrix, J., & Rocha, S. (2024). Navigating FRET Imaging Techniques for Biosensor Applications. 10.31219/OSF.IO/YE4DM

Cresens, C., Aytekin, S., Shaghaghi, B., Gerrits, L., Montero-Calle, A., Barderas, R., Kouwer, P., & Rocha, S. (2026). Protocol for synthesis and mechanical characterization of polyacrylamide hydrogels of varying stiffness for cell culture applications. STAR Protocols, 7(2), 104606. 10.1016/j.xpro.2026.104606

Crose, L. E. S., Hilder, T. L., Sciaky, N., & Johnson, G. L. (2009). Cerebral Cavernous Malformation 2 Protein Promotes Smad Ubiquitin Regulatory Factor 1-mediated RhoA Degradation in Endothelial Cells. The Journal of Biological Chemistry, 284(20), 13301. 10.1074/JBC.C900009200

Denier, C., Goutagny, S., Labauge, P., Krivosic, V., Arnoult, M., Cousin, A., Benabid, A. L., Comoy, J., Frerebeau, P., Gilbert, B., Houtteville, J. P., Jan, M., Lapierre, F., Loiseau, H., Menei, P., Mercier, P., Moreau, J. J., Nivelon-Chevallier, A., Parker, F.,… Tournier-Lasserve, E. (2004). Mutations within the MGC4607 Gene Cause Cerebral Cavernous Malformations. American Journal of Human Genetics, 74(2), 326–337. 10.1086/381718

Detter, M. R., Snellings, D. A., & Marchuk, D. A. (2018). Cerebral cavernous malformations develop through clonal expansion of mutant endothelial cells. Circulation Research, 123(10), 1143–1151. 10.1161/CIRCRESAHA.118.313970

Dulamea, A. O., & Lupescu, I. C. (2024). Cerebral cavernous malformations – An overview on genetics, clinical aspects and therapeutic strategies. Journal of the Neurological Sciences, 461. 10.1016/j.jns.2024.123044

Elosegui-Artola, A., Oria, R., Chen, Y., Kosmalska, A., Pérez-González, C., Castro, N., Zhu, C., Trepat, X., & Roca-Cusachs, P. (2016). Mechanical regulation of a molecular clutch defines force transmission and transduction in response to matrix rigidity. Nature Cell Biology, 18(5), 540–548. 10.1038/NCB3336

Faurobert, E., & Albiges-Rizo, C. (2010). Recent insights into cerebral cavernous malformations: A complex jigsaw puzzle under construction. FEBS Journal, 277(5), 1084–1096. 10.1111/J.1742-4658.2009.07537.X

Faurobert, E., Rome, C., Lisowska, J., Manet-Dupé, S., Boulday, G., Malbouyres, M., Balland, M., Bouin, A. P., Kéramidas, M., Bouvard, D., Coll, J. L., Ruggiero, F., Tournier-Lasserve, E., & Albiges-Rizo, C. (2013). CCM1–ICAP-1 complex controls β1 integrin–dependent endothelial contractility and fibronectin remodeling. The Journal of Cell Biology, 202(3), 545. 10.1083/JCB.201303044

Fernández-Martín, L., Marcos-Ramiro, B., Bigarella, C. L., Graupera, M., Cain, R. J., Reglero-Real, N., Jiménez, A., Cernuda-Morollón, E., Correas, I., Cox, S., Ridley, A. J., & Millán, J. (2012). Crosstalk between reticular adherens junctions and platelet endothelial cell adhesion molecule-1 regulates endothelial barrier function. Arteriosclerosis, Thrombosis, and Vascular Biology, 32(8). 10.1161/ATVBAHA.112.252080

Frischer, J. M., Pipp, I., Stavrou, I., Trattnig, S., Hainfellner, J. A., & Knosp, E. (2008). Cerebral cavernous malformations: congruency of histopathological features with the current clinical definition. Journal of Neurology, Neurosurgery & Psychiatry, 79(7), 783–788. 10.1136/JNNP.2007.132316

Gautam, U., Singh, M. B., Mahajan, S., Tripathi, S., Suri, V., Garg, A., Kumaran, S. S., Roy, S., & Suri, A. (2026). Biomechanical Characterization of Epileptic Brain for Surgical Planning of Intractable Epilepsy. Journal of Biomedical Materials Research Part A, 114(1), e70017. 10.1002/JBMA.70017

Glading, A., Han, J., Stockton, R. A., & Ginsberg, M. H. (2007). KRIT-1/CCM1 is a Rap1 effector that regulates endothelial cell–cell junctions. The Journal of Cell Biology, 179(2), 247. 10.1083/JCB.200705175

Goldstein, H. E., & Solomon, R. A. (2017). Epidemiology of cavernous malformations. Handbook of Clinical Neurology, 143, 241–247. 10.1016/B978-0-444-63640-9.00023-0

Grashoff, C., Hoffman, B. D., Brenner, M. D., Zhou, R., Parsons, M., Yang, M. T., McLean, M. A., Sligar, S. G., Chen, C. S., Ha, T., & Schwartz, M. A. (2010). Measuring mechanical tension across vinculin reveals regulation of focal adhesion dynamics. Nature, 466(7303), 263. 10.1038/NATURE09198

Hernandez-Varas, P., Berge, U., Lock, J. G., & Strömblad, S. (2015). A plastic relationship between vinculin-mediated tension and adhesion complex area defines adhesion size and lifetime. Nature Communications 2015 6:1, 6(1), 7524-. 10.1038/ncomms8524

Hong, C. C., Tang, A. T., Detter, M. R., Choi, J. P., Wang, R., Yang, X., Guerrero, A. A., Wittig, C. F., Hobson, N., Girard, R., Lightle, R., Moore, T., Shenkar, R., Polster, S. P., Goddard, L. M., Ren, A. A., Leu, N. A., Sterling, S., Yang, J.,… Kahn, M. L. (2020). Cerebral cavernous malformations are driven by ADAMTS5 proteolysis of versican. Journal of Experimental Medicine, 217(10). 10.1084/JEM.20200140/151938

Jilkova, Z. M., Lisowska, J., Manet, S., Verdier, C., Deplano, V., Geindreau, C., Faurobert, E., Albigès-Rizo, C., & Duperray, A. D. (2014). CCM proteins control endothelial β1 integrin dependent response to shear stress. Biology Open, 3(12), 1228–1235. 10.1242/BIO.201410132

Kanchanawong, P., & Calderwood, D. A. (2022). Organization, dynamics and mechanoregulation of integrin-mediated cell–ECM adhesions. Nature Reviews Molecular Cell Biology 2022 24:2, 24(2), 142–161. 10.1038/s41580-022-00531-5

Kiliç, T., Pamir, N., Küllü, S., Eren, F., Ozek, M. M., & Black, P. M. L. (2000). Expression of structural proteins and angiogenic factors in cerebrovascular anomalies. Neurosurgery, 46(5), 1179–1192. 10.1097/00006123-200005000-00032

Kim, D. H., & Wirtz, D. (2013). Focal adhesion size uniquely predicts cell migration. The FASEB Journal, 27(4), 1351. 10.1096/FJ.12-220160

Krishnan, R., Klumpers, D. D., Park, C. Y., Rajendran, K., Trepat, X., Van Bezu, J., Van Hinsbergh, V. W. M., Carman, C. V., Brain, J. D., Fredberg, J. J., Butler, J. P., & Van Nieuw Amerongen, G. P. (2011). Substrate stiffening promotes endothelial monolayer disruption through enhanced physical forces. American Journal of Physiology - Cell Physiology, 300(1), 146–154. 10.1152/ajpcell.00195.2010

Laberge-le Couteulx, S., Jung, H. H., Labauge, P., Houtteville, J. P., Lescoat, C., Cecillon, M., Marechal, E., Joutel, A., Bach, J. F., & Tournier-Lasserve, E. (1999). Truncating mutations in CCM1, encoding KRIT1, cause hereditary cavernous angiomas. Nature Genetics 1999 23:2, 23(2), 189–193. 10.1038/13815

LaCroix, A. S., Lynch, A. D., Berginski, M. E., & Hoffman, B. D. (2018). Tunable molecular tension sensors reveal extension-based control of vinculin loading. ELife, 7, e33927. 10.7554/ELIFE.33927

Lagendijk, A. K., Gomez, G. A., Baek, S., Hesselson, D., Hughes, W. E., Paterson, S., Conway, D. E., Belting, H. G., Affolter, M., Smith, K. A., Schwartz, M. A., Yap, A. S., & Hogan, B. M. (2017). Live imaging molecular changes in junctional tension upon VE-cadherin in zebrafish. Nature Communications 2017 8:1, 8(1), 1402-. 10.1038/s41467-017-01325-6

Lam, A. J., St-Pierre, F., Gong, Y., Marshall, J. D., Cranfill, P. J., Baird, M. A., McKeown, M. R., Wiedenmann, J., Davidson, M. W., Schnitzer, M. J., Tsien, R. Y., & Lin, M. Z. (2012). Improving FRET dynamic range with bright green and red fluorescent proteins. Nature Methods 2012 9:10, 9(10), 1005–1012. 10.1038/nmeth.2171

Liquori, C. L., Berg, M. J., Siegel, A. M., Huang, E., Zawistowski, J. S., Stoffer, T., Verlaan, D., Balogun, F., Hughes, L., Leedom, T. P., Plummer, N. W., Cannella, M., Maglione, V., Squitieri, F., Johnson, E. W., Rouleau, G. A., Ptacek, L., & Marchuk, D. A. (2003). Mutations in a Gene Encoding a Novel Protein Containing a Phosphotyrosine-Binding Domain Cause Type 2 Cerebral Cavernous Malformations. American Journal of Human Genetics, 73(6), 1459–1464. 10.1086/380314

Lisowska, J., Rödel, C. J., Manet, S., Miroshnikova, Y. A., Boyault, C., Planus, E., De Mets, R., Lee, H. H., Destaing, O., Mertani, H., Boulday, G., Tournier-Lasserve, E., Balland, M., Abdelilah-Seyfried, S., Albiges-Rizo, C., & Faurobert, E. (2018). The CCM1-CCM2 complex controls complementary functions of ROCK1 and ROCK2 that are required for endothelial integrity. Journal of Cell Science, 131(15). 10.1242/JCS.216093/56825

Malinverno, M., Maderna, C., Abu Taha, A., Corada, M., Orsenigo, F., Valentino, M., Pisati, F., Fusco, C., Graziano, P., Giannotta, M., Yu, Q. C., Zeng, Y. A., Lampugnani, M. G., Magnusson, P. U., & Dejana, E. (2019). Endothelial cell clonal expansion in the development of cerebral cavernous malformations. Nature Communications 2019 10:1, 10(1), 2761-. 10.1038/s41467-019-10707-x

Maruthamuthu, V., Sabass, B., Schwarz, U. S., & Gardel, M. L. (2011). Cell-ECM traction force modulates endogenous tension at cell-cell contacts. Proceedings of the National Academy of Sciences of the United States of America, 108(12), 4708–4713. 10.1073/PNAS.1011123108

Millon-Frémillon, A., Bouvard, D., Grichine, A., Manet-Dupé, S., Block, M. R., & Albiges-Rizo, C. (2008). Cell adaptive response to extracellular matrix density is controlled by ICAP-1–dependent β1-integrin affinity. Journal of Cell Biology, 180(2), 427–441. 10.1083/JCB.200707142

Morsut, L., Roybal, K. T., Xiong, X., Gordley, R. M., Coyle, S. M., Thomson, M., & Lim, W. A. (2016). Engineering Customized Cell Sensing and Response Behaviors Using Synthetic Notch Receptors. Cell, 164(4), 780. 10.1016/J.CELL.2016.01.012

Pannekoek, W. J., Post, A., & Bos, J. L. (2014). Rap1 signaling in endothelial barrier control. Cell Adhesion & Migration, 8(2), 100. 10.4161/CAM.27352

Pelham, R. J., & Wang, Y. L. (1997). Cell locomotion and focal adhesions are regulated by substrate flexibility. Proceedings of the National Academy of Sciences of the United States of America, 94(25), 13661–13665. 10.1073/PNAS.94.25.13661

Postma, R. J., Fischer, S. E., Bijkerk, R., & van Zonneveld, A. J. (2025). Unveiling endothelial cell border heterogeneity: VE-cadherin adherens junction stratification by deep convolutional neural networks. PLOS ONE, 20(1), e0317110. 10.1371/JOURNAL.PONE.0317110

Ranjit, S., Malacrida, L., Jameson, D. M., & Gratton, E. (2018). Fit-free analysis of fluorescence lifetime imaging data using the phasor approach. Nature Protocols 2018 13:9, 13(9), 1979–2004. 10.1038/s41596-018-0026-5

Rath, M., Pagenstecher, A., Hoischen, A., & Felbor, U. (2020). Postzygotic mosaicism in cerebral cavernous malformation. Journal of Medical Genetics, 57(3), 212–216.10.1136/JMEDGENET-2019-106182

Ren, J., Wang, D., Wang, L., Jiang, C., Tian, A., Cui, Z., Ren, Y., Bian, L., Zeng, G., Meng, G., Shan, Y., Liang, J., Xiao, X., Tang, J., Wei, Y., He, C., Sun, L., Ma, Y., Yu, J.,… Hong, T. (2025). Clinical, genomic, and histopathologic diversity in cerebral cavernous malformations. Acta Neuropathologica Communications 2025 13:1, 13(1), 23-.10.1186/S40478-025-01940-1

Ringer, P., Weißl, A., Cost, A. L., Freikamp, A., Sabass, B., Mehlich, A., Tramier, M., Rief, M., & Grashoff, C. (2017). Multiplexing molecular tension sensors reveals piconewton force gradient across talin-1. Nature Methods 2017 14:11, 14(11), 1090–1096. 10.1038/nmeth.4431

Rödel, C. J., & Abdelilah-Seyfried, S. (2026). CCM scaffolds at the frontier of biomechanics: their roles in vascular physiology and disease. Biological Chemistry, 407(4–6), 247–258. 10.1515/HSZ-2025-0246/XML

Rothbart, D., Awad, I. A., Lee, J., Kim, J., Harbaugh, R., & Criscuolo, G. R. (1996). Expression of angiogenic factors and structural proteins in central nervous system vascular malformations. Neurosurgery, 38(5), 915–925. 10.1097/00006123-199605000-00011

Rothenberg, K. E., Scott, D. W., Christoforou, N., & Hoffman, B. D. (2018). Vinculin Force-Sensitive Dynamics at Focal Adhesions Enable Effective Directed Cell Migration. Biophysical Journal, 114(7), 1680. 10.1016/J.BPJ.2018.02.019

Sarangi, B. R., Gupta, M., Doss, B. L., Tissot, N., Lam, F., Mège, R. M., Borghi, N., & Ladoux, B. (2017). Coordination between Intra- and Extracellular Forces Regulates Focal Adhesion Dynamics. Nano Letters, 17(1), 399–406. 10.1021/ACS.NANOLETT.6B04364

Shapeti, A., Barrasa-Fano, J., Abdel Fattah, A. R., de Jong, J., Sanz-Herrera, J. A., Pezet, M., Assou, S., de Vet, E., Elahi, S. A., Ranga, A., Faurobert, E., & Van Oosterwyck, H. (2024). Force-mediated recruitment and reprogramming of healthy endothelial cells drive vascular lesion growth. Nature Communications 2024 15:1, 15(1), 8660-.10.1038/s41467-024-52866-6

Shoyer, T. C., Gates, E. M., Cabe, J. I., Urs, A. N., Conway, D. E., & Hoffman, B. D. (2023). Coupling during collective cell migration is controlled by a vinculin mechanochemical switch. Proceedings of the National Academy of Sciences of the United States of America, 120(50), e2316456120. 10.1073/PNAS.2316456120

Snellings, D. A., Hong, C. C., Ren, A. A., Lopez-Ramirez, M. A., Girard, R., Srinath, A., Marchuk, D. A., Ginsberg, M. H., Awad, I. A., & Kahn, M. L. (2021). Cerebral Cavernous Malformation: From Mechanism to Therapy. Circulation Research, 129(1), 195–215. 10.1161/CIRCRESAHA.121.318174

Stockton, R. A., Shenkar, R., Awad, I. A., & Ginsberg, M. H. (2010). Cerebral cavernous malformations proteins inhibit Rho kinase to stabilize vascular integrity. The Journal of Experimental Medicine, 207(4), 881. 10.1084/JEM.20091258

Swamy, H., & Glading, A. J. (2022). Is Location Everything? Regulation of the Endothelial CCM Signaling Complex. Frontiers in Cardiovascular Medicine, 9, 954780. 10.3389/FCVM.2022.954780

Tan, J. L., Tien, J., Pirone, D. M., Gray, D. S., Bhadriraju, K., & Chen, C. S. (2003). Cells lying on a bed of microneedles: An approach to isolate mechanical force. Proceedings of the National Academy of Sciences, 100(4), 1484–1489. 10.1073/PNAS.0235407100

Urbano, R. L., Furia, C., Basehore, S., & Clyne, A. M. (2017). Stiff Substrates Increase Inflammation-Induced Endothelial Monolayer Tension and Permeability. Biophysical Journal, 113(3), 645–655. 10.1016/j.bpj.2017.06.033

Urbanski, M. M., Brendel, M. B., & Melendez-Vasquez, C. V. (2019). Acute and chronic demyelinated CNS lesions exhibit opposite elastic properties. Scientific Reports 2019 9:1, 9(1), 999-. 10.1038/s41598-018-37745-7

Uttekar, N., Artz, A., Ghanekar, V., Kaul, P., Heinrichs, J., Stegmeyer, R. I., Takir, G. G., Nottebaum, A. F., & Vestweber, D. (2026). Plakoglobin transmits tension across VE-cadherin for vascular leak formation and leukocyte diapedesis. The EMBO Journal, 45(7), 2210. 10.1038/S44318-026-00732-0

Vannier, D. R., Shapeti, A., Chuffart, F., Planus, E., Manet, S., Rivier, P., Destaing, O., Albiges-Rizo, C., Van Oosterwyck, H., & Faurobert, E. (2021). CCM2-deficient endothelial cells undergo a ROCK-dependent reprogramming into senescence-associated secretory phenotype. Angiogenesis 2021 24:4, 24(4), 843–860. 10.1007/S10456-021-09809-2

Voss, K., Stahl, S., Schleider, E., Ullrich, S., Nickel, J., Mueller, T. D., & Felbor, U. (2007). CCM3 interacts with CCM2 indicating common pathogenesis for cerebral cavernous malformations. Neurogenetics 2007 8:4, 8(4), 249–256. 10.1007/S10048-007-0098-9

Whitehead, K. J., Chan, A. C., Navankasattusas, S., Koh, W., London, N. R., Ling, J., Mayo, A. H., Drakos, S. G., Marchuk, D. A., Davis, G. E., & Li, D. Y. (2009). The Cerebral Cavernous Malformation signaling pathway promotes vascular integrity via Rho GTPases. Nature Medicine, 15(2), 177. 10.1038/NM.1911

Wong, J. H., Awad, I. A., & Kim, J. H. (2000). Ultrastructural pathological features of cerebrovascular malformations: a preliminary report. Neurosurgery, 46(6), 1454–1459. 10.1097/00006123-200006000-00027

Yang, D., Tao, Z., Chen, S., Wang, H., Luo, L., & Chen, J. (2026). Loss of arap3 drives a cerebral cavernous malformation-like phenotype in zebrafish via Krit1 interaction and RhoA dysregulation. Development (Cambridge, England), 153(18). 10.1242/DEV.205762/372968

Yang, X., Zhang, J., Dai, Z., Yang, T., Xie, L., Gao, F., Zheng, X., & Han, Z. (2026). Knockin expression of human ADAMTS5 impairs cardiovascular development and aggravates cerebral cavernous malformations in mice. DMM Disease Models and Mechanisms, 19(5). 10.1242/DMM.052668/371848

Yin, J., Maggi, L., Wiesner, C., Affolter, M., & Belting, H. G. (2024). Oscillatory contractile forces refine endothelial cell-cell interactions for continuous lumen formation governed by Heg1/Ccm1. Angiogenesis 2024 27:4, 27(4), 845–860. 10.1007/S10456-024-09945-5

Zhou, Z., Rawnsley, D. R., Goddard, L. M., Pan, W., Cao, X. J., Jakus, Z., Zheng, H., Yang, J., Arthur, J. S. C., Whitehead, K. J., Li, D., Zhou, B., Garcia, B. A., Zheng, X., & Kahn, M. L. (2015). The Cerebral Cavernous Malformation Pathway Controls Cardiac Development via Regulation of Endocardial MEKK3 Signaling and KLF Expression. Developmental Cell, 32(2), 168–180. 10.1016/j.devcel.2014.12.009

