## Supplementary Figures for "Letting go to hold on: loss of CCM2 disrupts endothelial force balance in cerebral cavernous malformations"

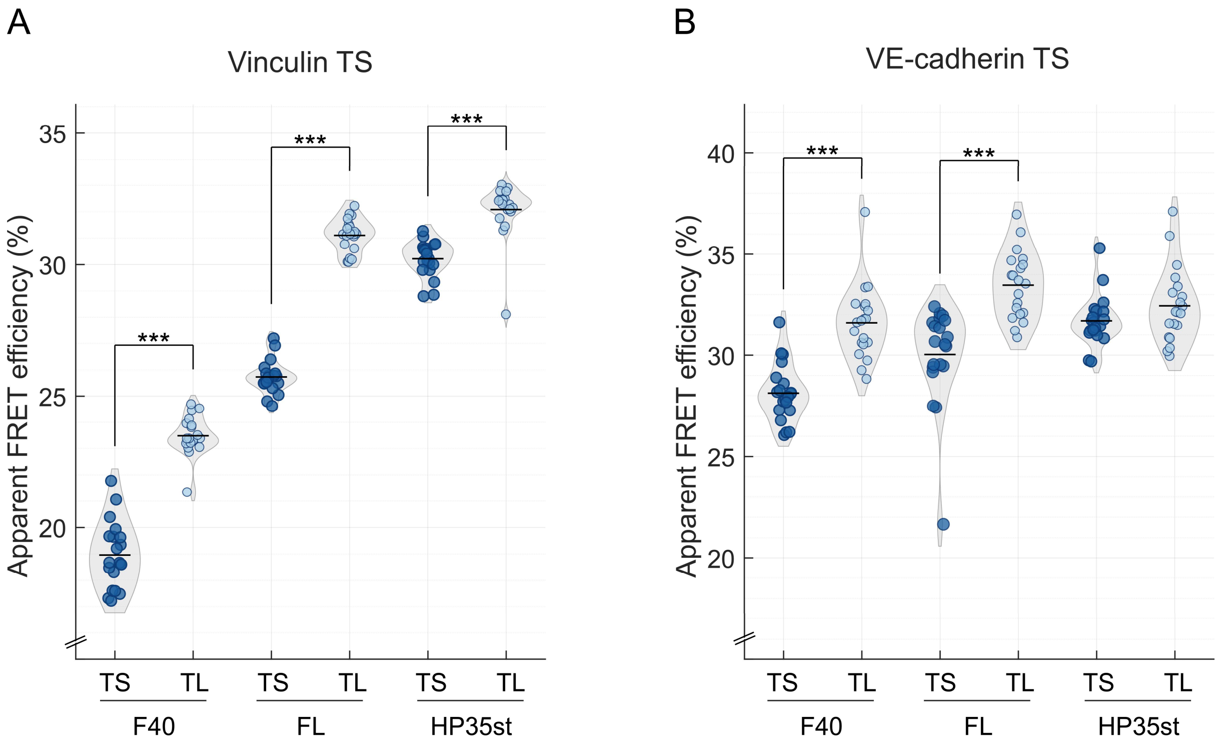


| Condition | Mean | SD | Δ |
|---|---|---|---|
| TS_F40 | 28.135 | ±1.42 | 3.47 % |
| TL_F40 | 31.609 | ±1.81 | |
| TS_FL | 30.033 | ±2.44 | 3.43 % |
| TL_FL | 33.466 | ±1.65 | |
| TS_HP35st | 31.705 | ±1.24 | 0.742 % |
| TL_HP35st | 32.447 | ±1.85 | |

| Condition | Mean | SD | Δ |
|---|---|---|---|
| TS_F40 | 18.96 | ±1.25 | 4.53 % |
| TL_F40 | 23.49 | ±0.73 | |
| TS_FL | 25.727 | ±0.64 | 5.39 % |
| TL_FL | 31.115 | ±0.62 | |
| TS_HP35st | 30.222 | ±0.66 | 1.87 % |
| TL_HP35st | 32.092 | ±1.03 | |

**Supplementary Figure 1: Selection of elastic linker for (A) vinculin and (B) VE-cadherin tension sensor.** The graphs show violin scatter plots of FRET efficiency indicative of tension load between each linker (F40/FL/HP35st) within the full tension sensor (TS – dark blue) and the tailless (TL – light blue) control. Statistical analysis between TS and TL per linker was performed using paired t-tests (see Methods). The tables display the exact mean values with the standard deviation (SD) and the difference between TS and TL per linker with the best separation highlighted in green. N = 18-20 per violin plot, *p < 0.05, **p < 0.01, ***p < 0.001.

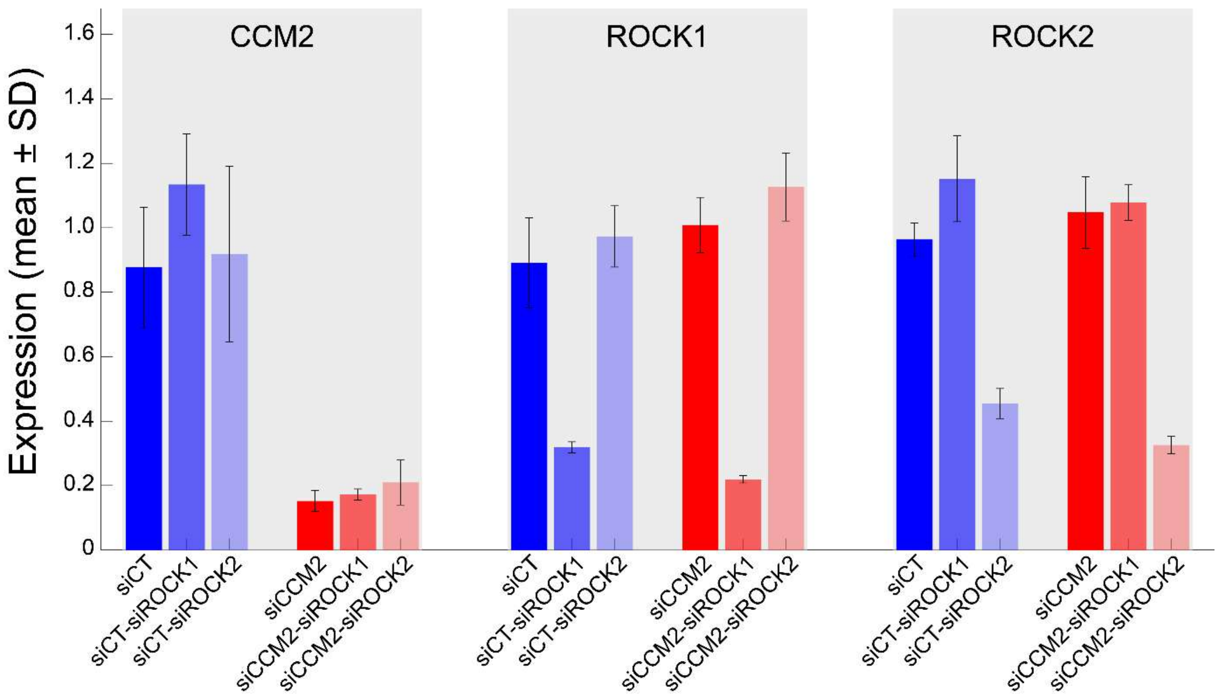


**Supplementary Figure 2: siRNA silencing efficiency in HBMVECs.** qPCR analysis of CCM2, ROCK1 and ROCK2 mRNA expression levels normalized to siCT in molecular tension HBMVEC cell lines following siRNA silencing. N = 4-6 from 2 biological replicates.

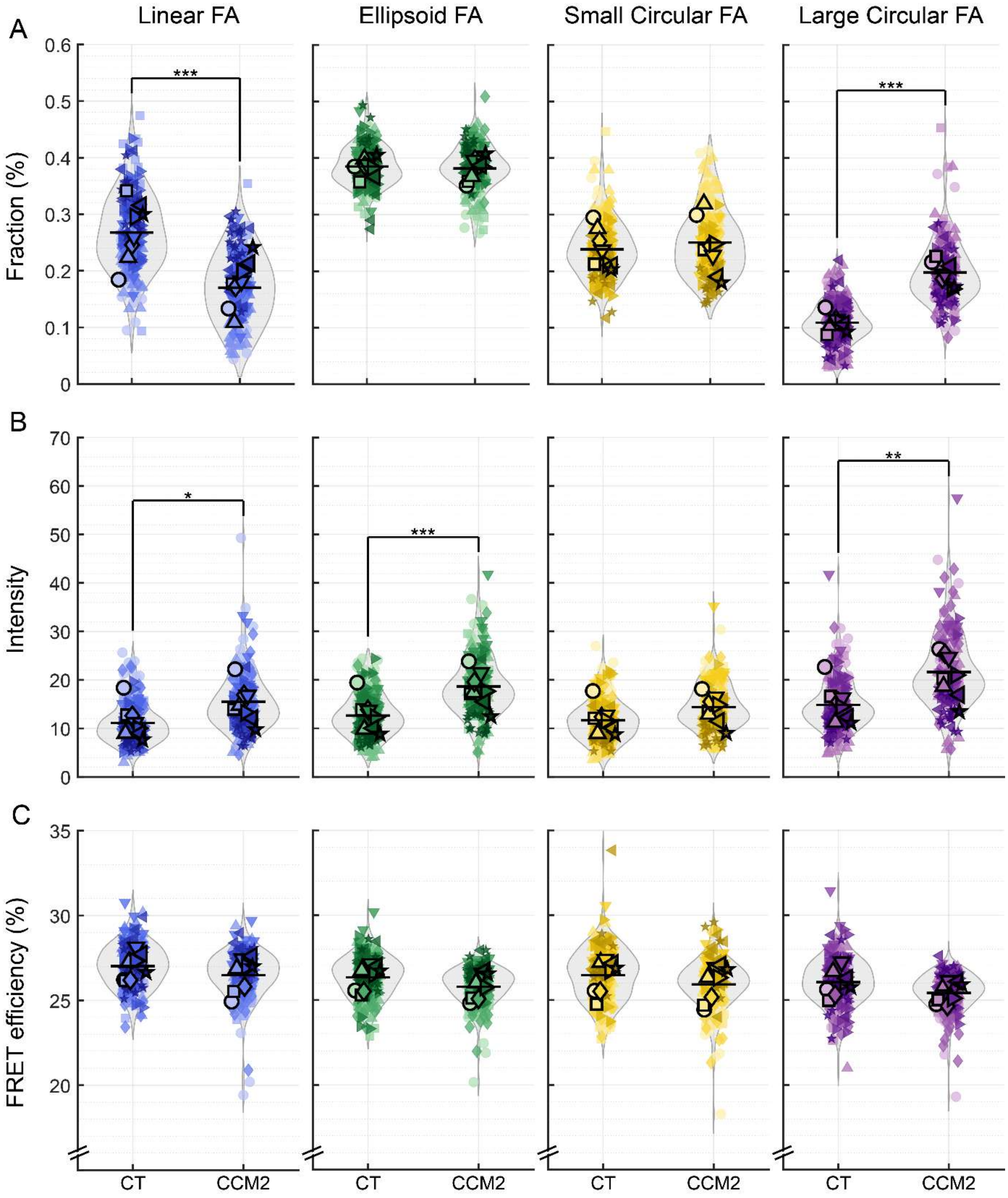


**Supplementary Figure 3: Comparison of CT vs CCM2 on glass separated per morphological FA category.** The average (A) fraction, (B) intensity and (C) FRET efficiency of siCT and siCCM2 samples compared per category: linear (blue), ellipsoid (green), small (yellow) and large circular (purple) FAs. Statistical analysis between all groups was performed using linear-mixed effects model on the whole dataset (see Methods). N = 217 for CT and 183 for CCM2 from 8 biological replicates, *$p < 0.05$, **$p < 0.01$, ***$p < 0.001$.

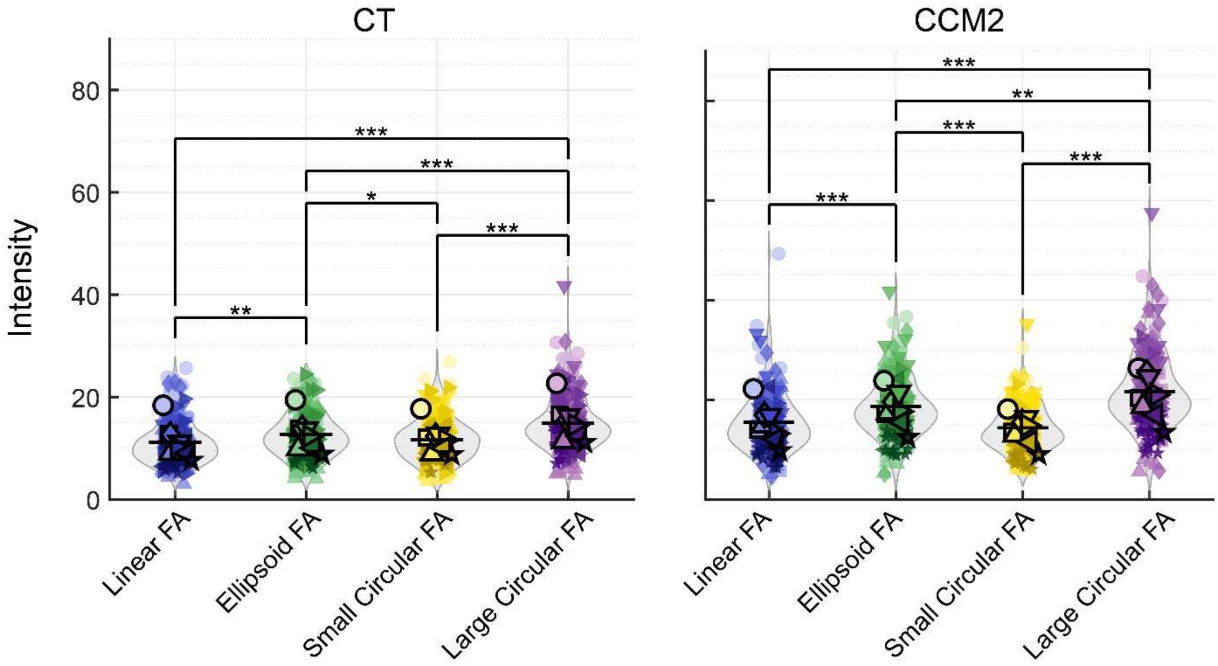


**Supplementary Figure 4: Comparison of average intensity values between morphological FA categories divided between CT or CCM2 condition.** Statistical analysis between all groups was performed using repeat measures ANOVA within each condition (see Methods). N = 217 for CT and 183 for CCM2 from 8 biological replicates for each morphology, *p < 0.05, **p < 0.01, ***p < 0.001.

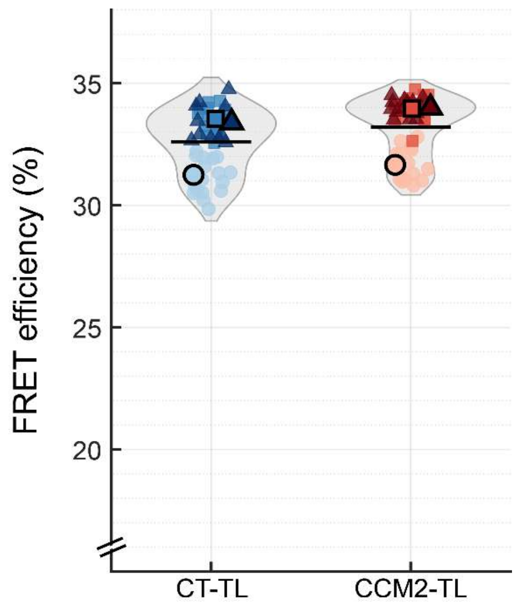


**Supplementary Figure 5: Vinculin tension-insensitive control (TL) under CT and CCM2-depleted conditions.** CCM2 silencing did not result in a change in FRET efficiency compared to its CT control. Statistically analyzed using independent paired t-test (see Methods). N = 49 for CT-TL and 45 for CCM2-TL from 3 biological replicates, *p < 0.05, **p < 0.01, ***p < 0.001.

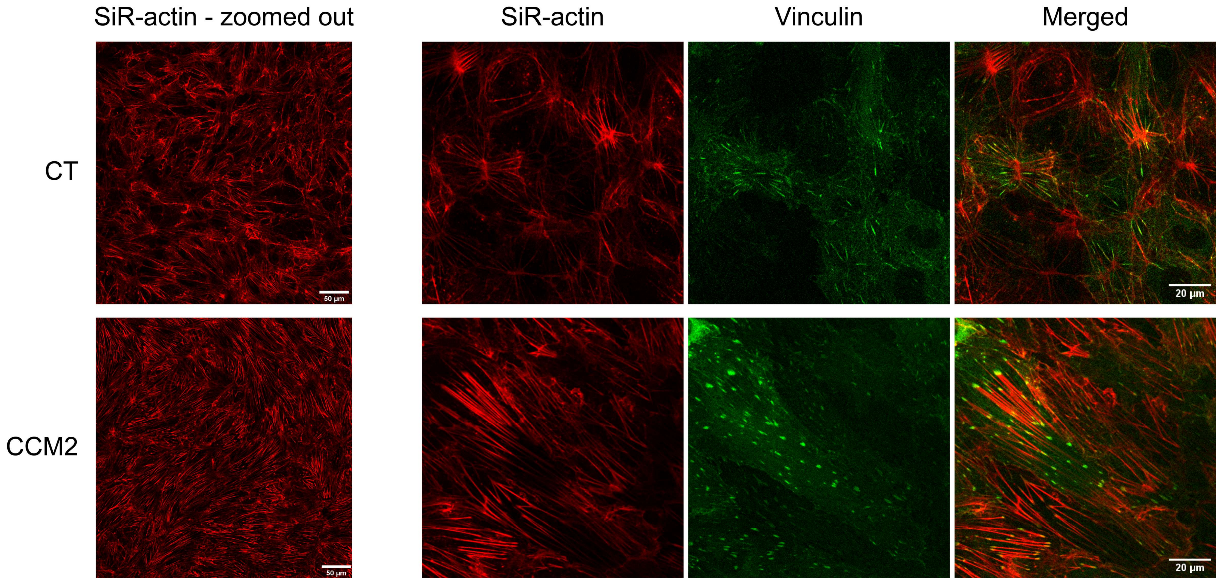


**Supplementary Figure 6: Actin staining and vinculin in CT and CCM2 depleted HBMVECs.** CCM2 depleted monolayers show an increase in stress fibers which attach to more prevalent, larger and more dens FAs. Left: SiR-actin shows the stained actin network in HBMVEC monolayers. Right: Overlay of SiR-actin with vinculin visualized through expression of VinTS tension sensor in the HBMVEC monolayer.

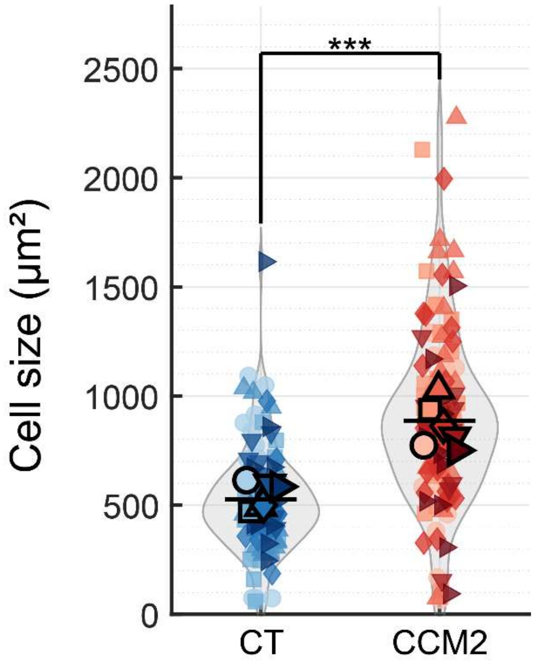


**Supplementary Figure 7: Comparison average cell size between healthy (CT) and CCM2-depleted HBMVEC monolayers.** N = 123 for CT and 133 for CCM2 from 6 biological replicates, *$p < 0.05$, **$p < 0.01$, ***$p < 0.001$. This number is lower compared to other parameters calculated due to removal of FOV where segmentation was not sufficient.

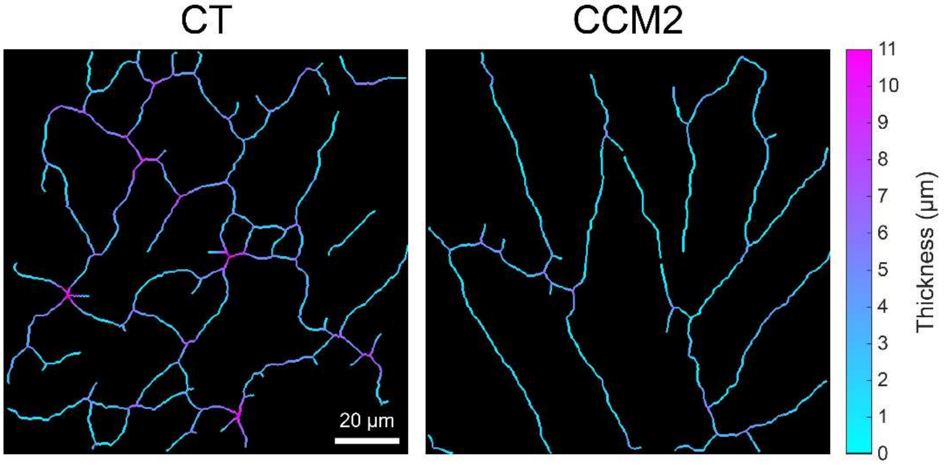


**Supplementary Figure 8: Representation of adherens junction thickness along the VE-cadherin skeleton.** Medial-axis skeletons of the junctional network, with each skeleton pixel colored by the local junction thickness (µm) according to the color scale shown on the right side. Thickness was obtained from a Euclidean distance transform of the smoothed mask, in which each skeleton pixel takes the diameter of the largest disk that fits inside the junction at that position (see Methods).

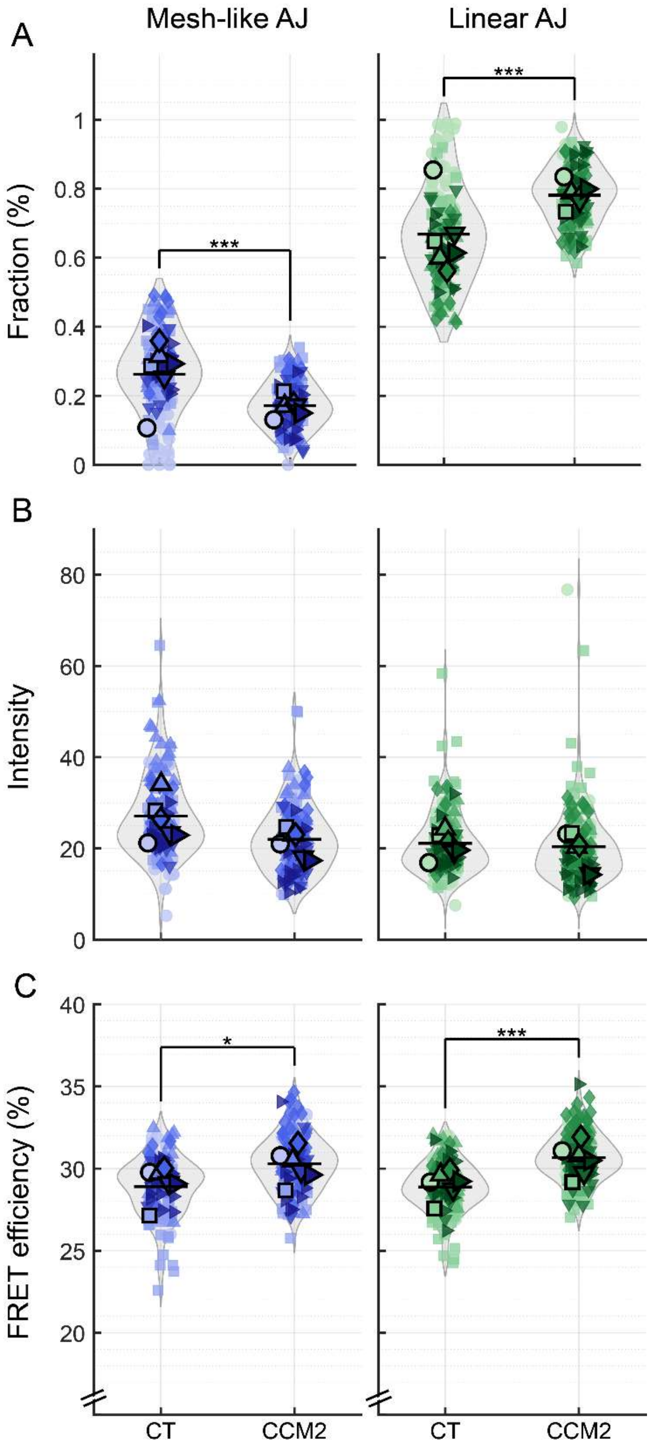


**Supplementary Figure 9: Comparison of CT vs CCM2 on glass separated per morphological AJ category.** The average (A) fraction, (B) intensity and (C) FRET efficiency of siCT and siCCM2 samples compared per category: mesh-like (blue) and linear (green) AJs. Statistical analysis between all groups was performed using linear-mixed effects model on the whole dataset (see Methods). N = 140 for CT and 154 for CCM2 from 6 biological replicates, *$p < 0.05$, **$p < 0.01$, ***$p < 0.001$.

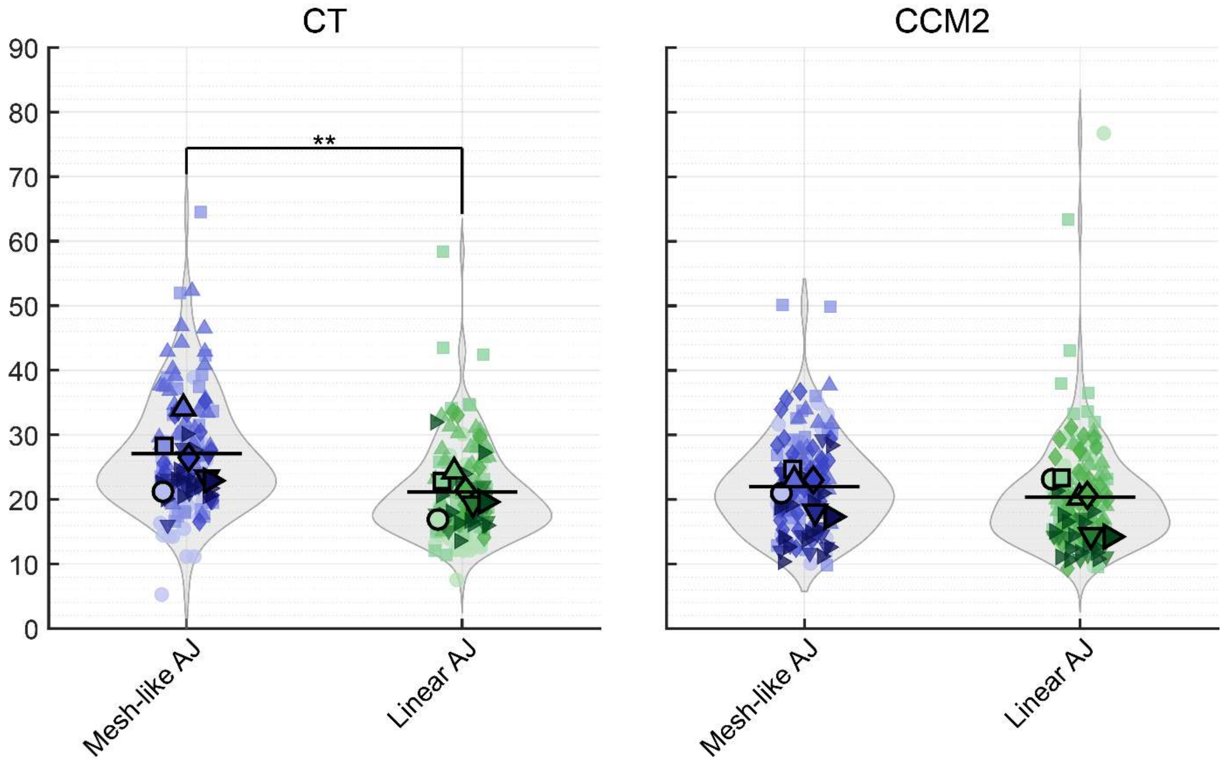


**Supplementary Figure 10: Comparison of average intensity values between morphological AJ categories divided between CT or CCM2 condition.** Statistical analysis between all groups was performed using repeat measures ANOVA within each condition (see Methods). N = 140 for CT and 154 for CCM2 from 6 biological replicates for each morphology, $*p < 0.05$, $**p < 0.01$, $***p < 0.001$.

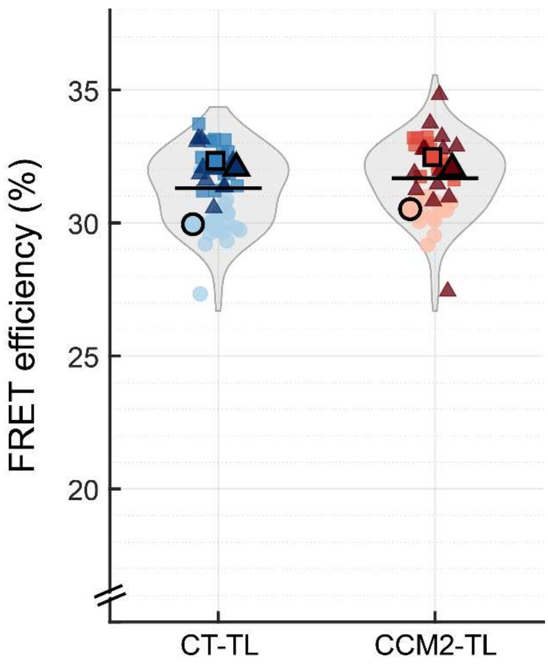


**Supplementary Figure 11: VE-cadherin tension-insensitive control (TL) under CT and CCM2-depleted conditions.** CCM2 silencing did not result in a change in FRET efficiency compared to its CT control. Statistically analyzed using independent paired t-test (see Methods). N = 50 for CT-TL and 45 for CCM2-TL from 3 biological replicates, *$p < 0.05$, **$p < 0.01$, ***$p < 0.001$.

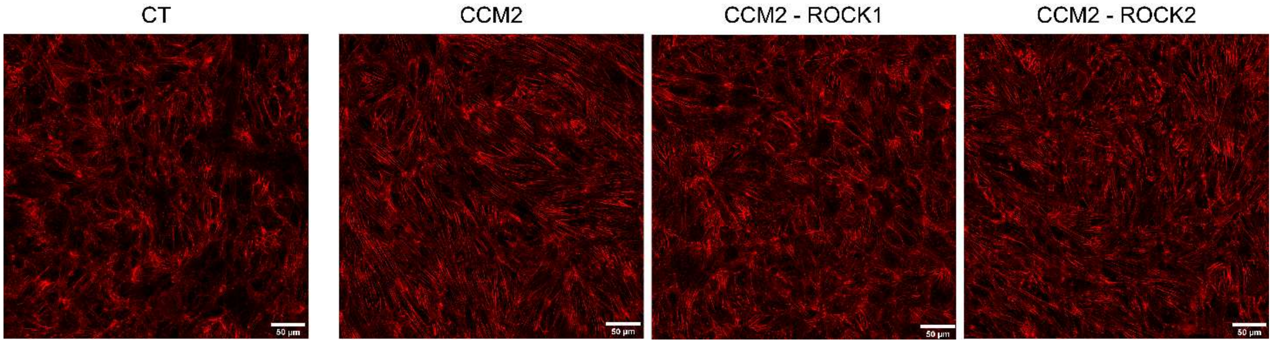


**Supplementary Figure 12: Actin staining in CT, CCM2, CCM2-ROCK1 and CCM2-ROCK2 depleted HBMVECs.** SiR-actin stained HBMVEC monolayers show increased stress fibers in CCM2 samples, which get decreased toward healthy (CT) monolayers more so in CCM2-ROCK1 samples than in CCM2-ROCK2 samples.

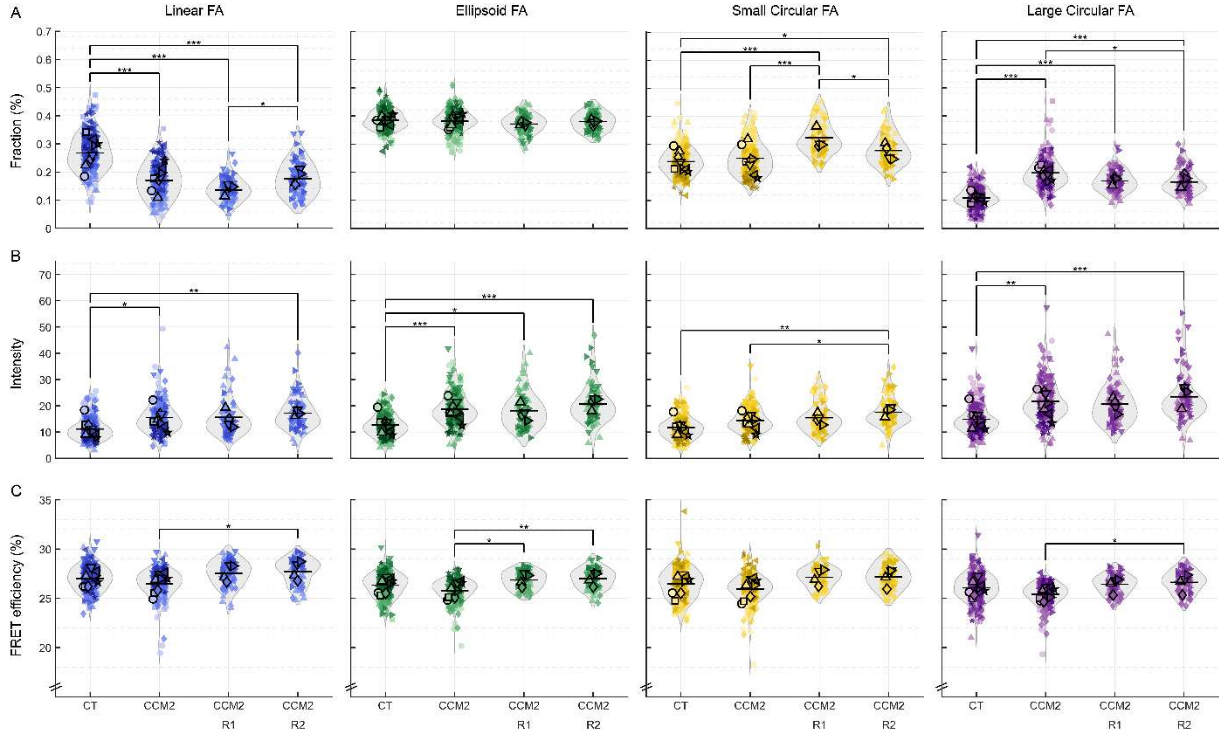


**Supplementary Figure 13: Comparison of CT, CCM2 and CCM2 with ROCK silencing on glass separated per morphological FA category.** The average (A) fraction, (B) intensity and (C) FRET efficiency of siCT, siCCM2, siCCM2-siROCK1, siCCM2-siROCK2 samples compared per category: linear (blue), ellipsoid (green), small (yellow) and large circular (purple) FAs. Statistical analysis between all groups was performed using linear-mixed effects model on the whole dataset (see Methods). N = 217, 183, 73, 73 for siCT, siCCM2, siCCM2-siROCK1, siCCM2-siROCK2 respectively from 8 or 4 biological replicates, *p < 0.05, **p < 0.01, ***p < 0.001.

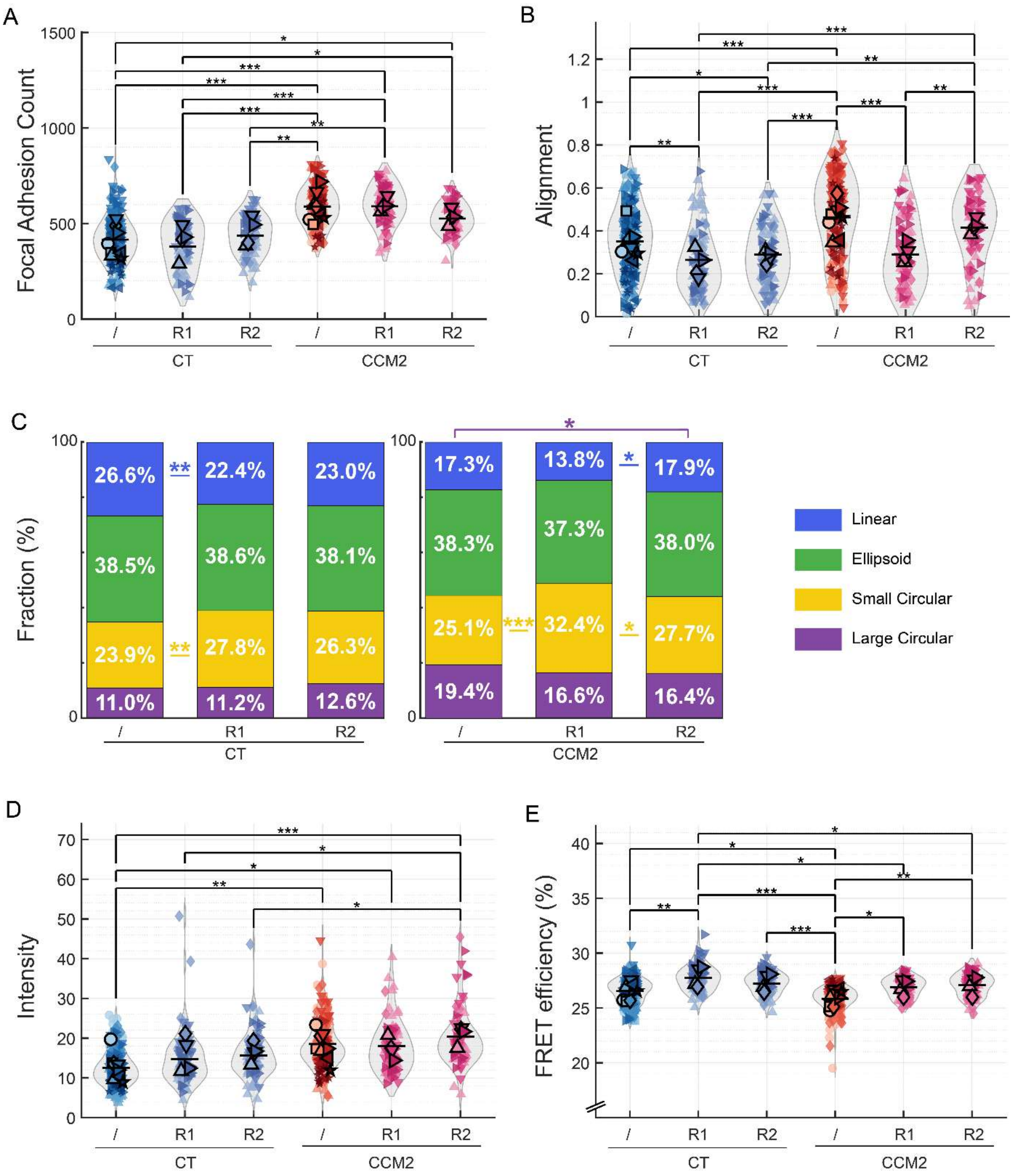


**Supplementary Figure 14: Effect of ROCK silencing on FA in healthy (CT) and CCM2-depleted monolayers.** (A) FA count per FOV. (B) FA alignment within a given FOV (non-circular FAs only). (C) Stacked histogram representing the fraction of each morphological group per condition. (D) Average vinculin intensity representing vinculin recruitment. (E) Average FRET efficiency indicative of tension load. Statistical analysis between all groups was performed using linear-mixed effects model on the whole dataset (see Methods). N = 217, 85, 82, 183, 73, 73 for siCT, siCT-siROCK1, siCT-siROCK2, siCCM2, siCCM2-siROCK1, siCCM2-siROCK2 respectively from 8 or 4 biological replicates,*p < 0.05, **p < 0.01, ***p < 0.001.

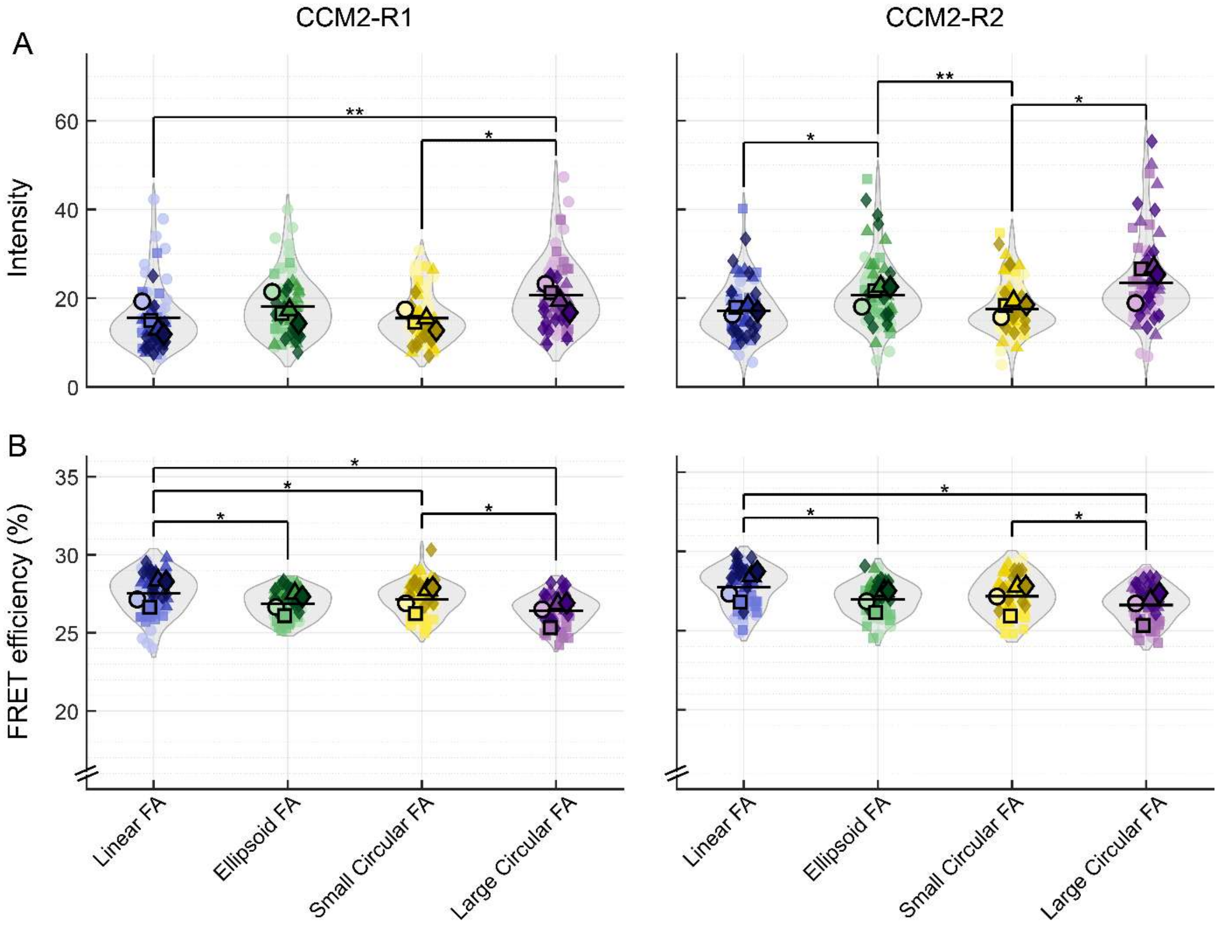


**Supplementary Figure 15: Comparison between morphological FA categories separated by CCM2-ROCK1 and CCM2-ROCK2 conditions.** The average (A) intensity and (B) FRET efficiency of siCCM2-siROCK1 and siCCM2-siROCK2 further separated by FA category. (Values for siCT and siCCM2 can be found in main figure 2G and supplementary figure 4) N = 73 and 73 for siCCM2-siROCK1 and siCCM2-siROCK2 respectively from 4 biological replicates, *$p < 0.05$, **$p < 0.01$, ***$p < 0.001$.

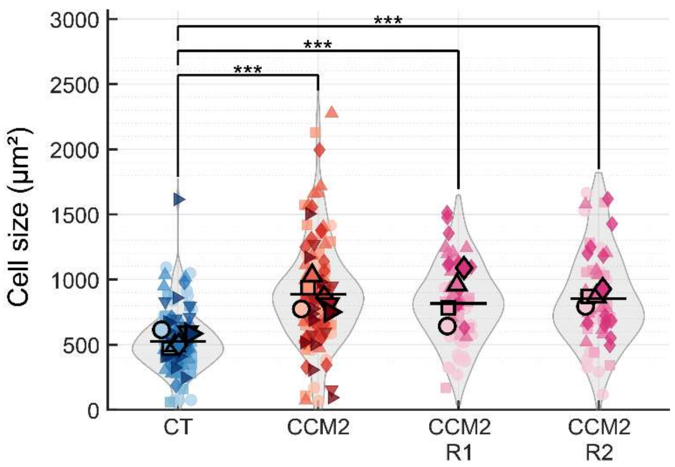


**Supplementary Figure 16: Comparison average cell size between CT, CCM2 and CCM2 with ROCK depleted HBMVEC monolayers.** N = 123, 133, 65, 63 for siCT, siCCM2, siCCM2-siROCK1, siCCM2-siROCK2 respectively from 6 or 4 biological replicates, $*p < 0.05$, $**p < 0.01$, $***p < 0.001$. This number is lower compared to other parameters calculated due to removal of FOV where segmentation was not sufficient.

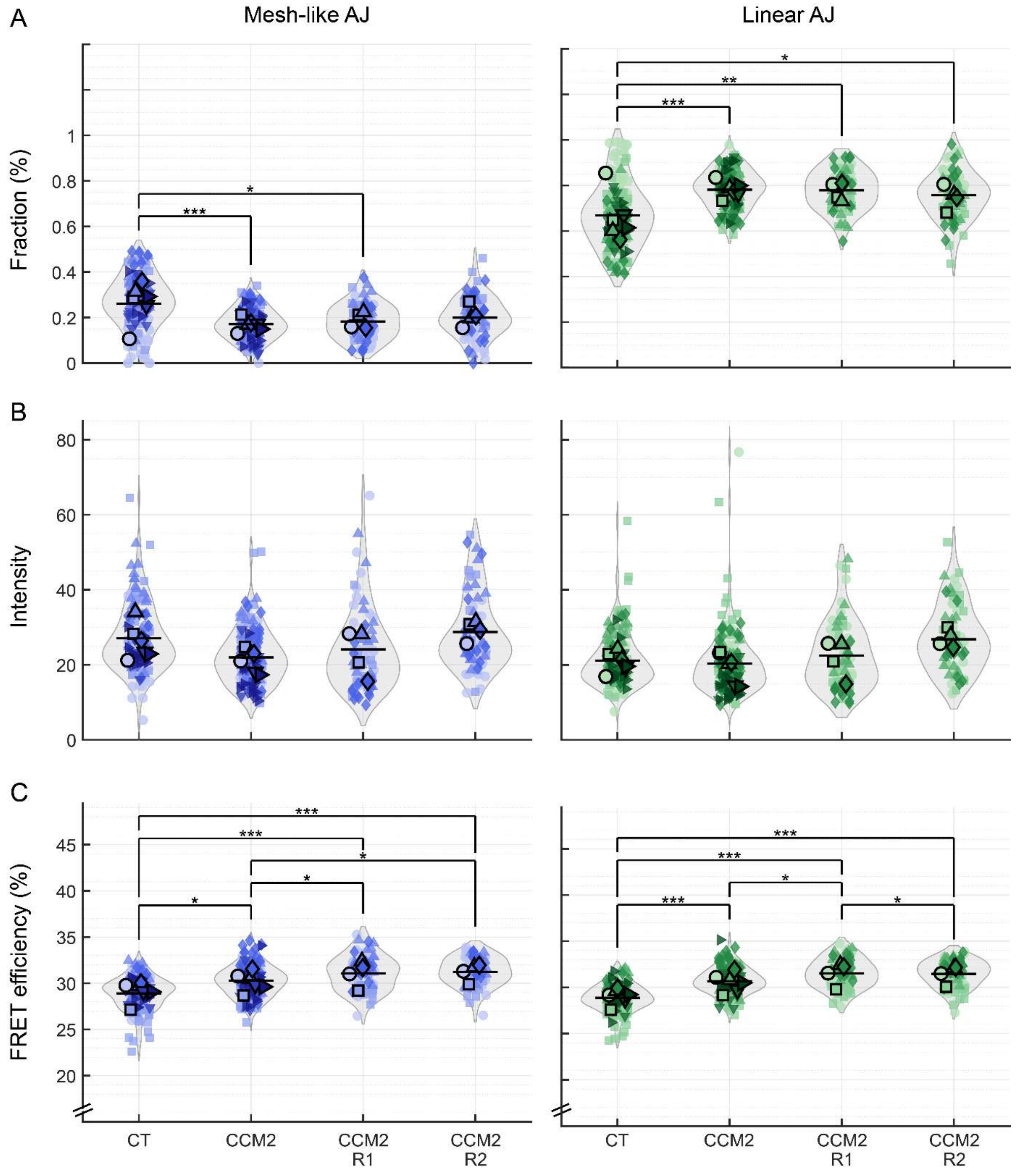


**Supplementary Figure 17: Comparison of CT, CCM2 and CCM2 with ROCK silencing on glass separated per morphological AJ category.** The average (A) fraction, (B) intensity and (C) FRET efficiency of siCT, siCCM2, siCCM2-siROCK1, siCCM2-siROCK2 samples compared per category: mesh-like (blue) and linear (green) AJs. Statistical analysis between all groups was performed using linear-mixed effects model on the whole dataset (see Methods). N = 140, 154, 73, 72 for siCT, siCCM2, siCCM2-siROCK1, siCCM2-siROCK2 respectively from 6 or 4 biological replicates, $*p < 0.05$, $**p < 0.01$, $***p < 0.001$.

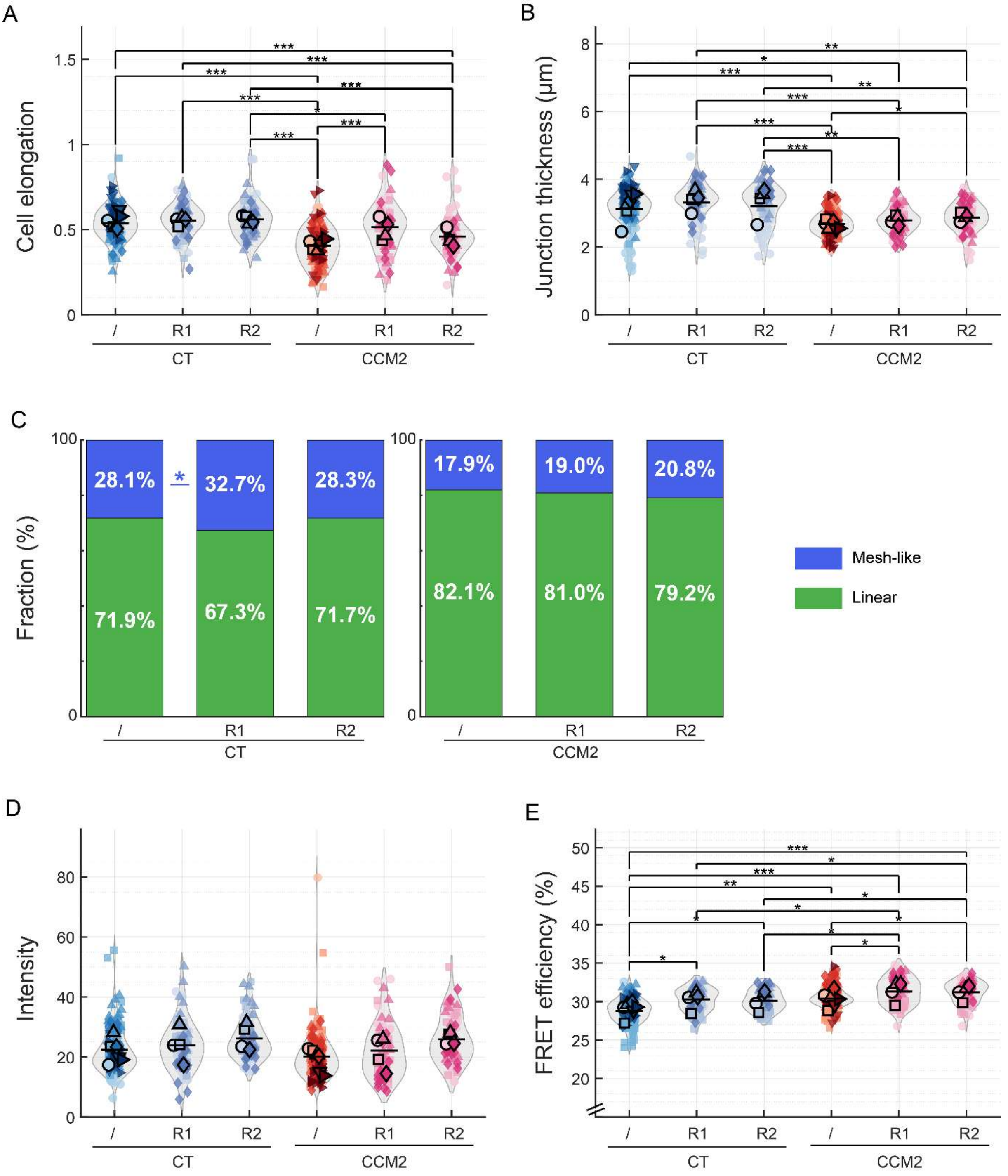


**Supplementary Figure 18: Effect of ROCK silencing on AJ in healthy (CT) and CCM2-depleted monolayers.** (A) Average cell elongation calculated through the major and minor axis ratio of every cell. (B) Average AJ thickness. (C) Stacked histogram representing the fraction of each morphological group per condition. (D) Average VE-cadherin intensity representing VE-cadherin recruitment. (E) Average FRET efficiency indicative of tension load. Statistical analysis between all groups was performed using linear-mixed effects model on the whole dataset (see Methods). N = 140, 74, 73, 154, 73, 72 for siCT, siCT-siROCK1, siCT-siROCK2, siCCM2, siCCM2-siROCK1, siCCM2-siROCK2 respectively from 6 or 4 biological replicates, $*p < 0.05$, $**p < 0.01$, $***p < 0.001$.

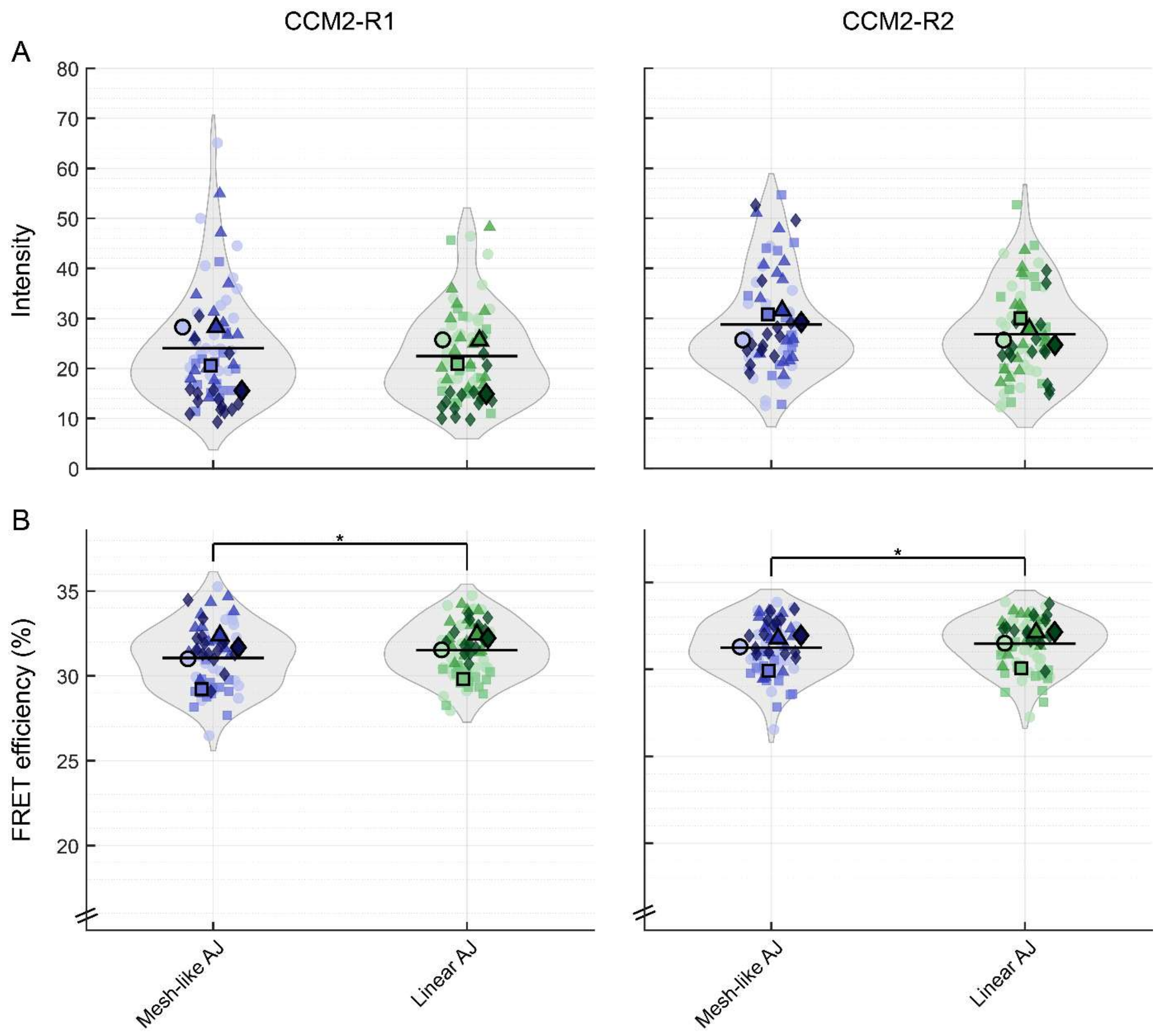


**Supplementary Figure 19: Comparison between morphological AJ categories separated by CCM2-ROCK1 and CCM2-ROCK2 conditions.** The average (A) intensity and (B) FRET efficiency of siCCM2-siROCK1 and siCCM2-siROCK2 further separated by AJ category. (Values for siCT and siCCM2 can be found in main figure 2N and supplementary figure 10) N = 73 and 72 for siCCM2-siROCK1 and siCCM2-siROCK2 respectively from 4 biological replicates, *$p < 0.05$, **$p < 0.01$, ***$p < 0.001$.

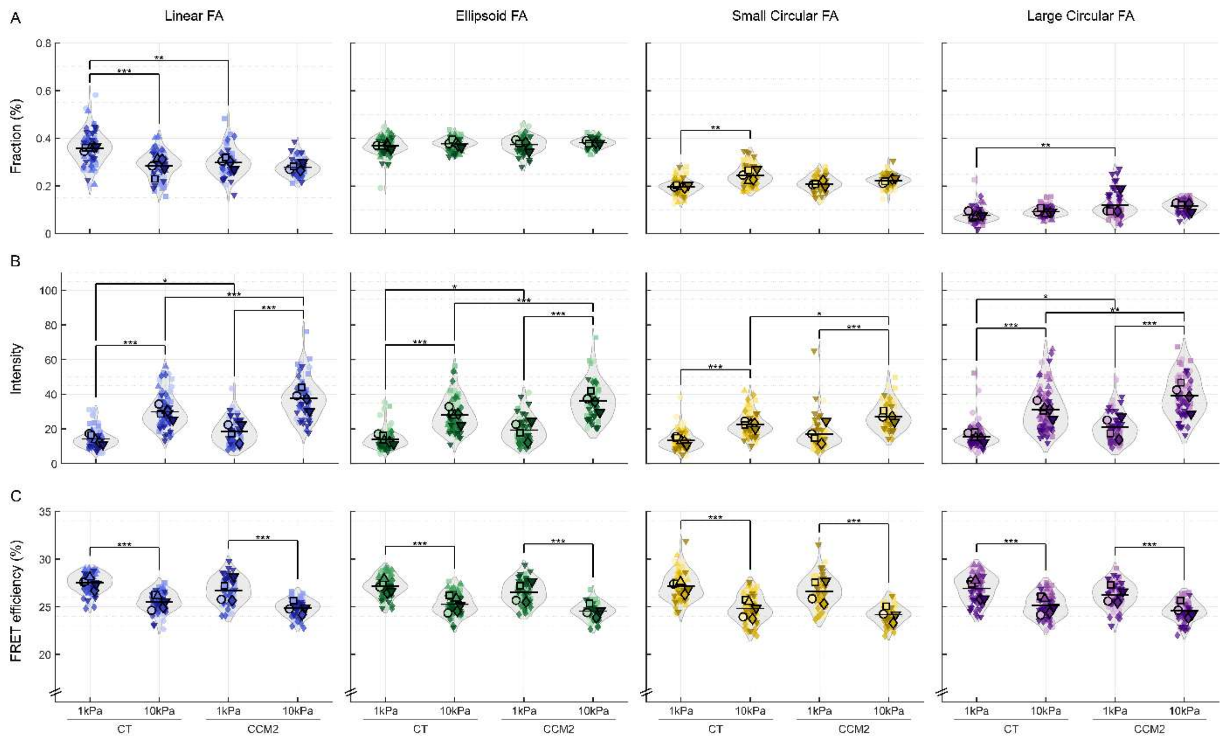


**Supplementary Figure 20: Comparison of CT and CCM2 on gel per morphological FA category.** The average (A) fraction, (B) intensity and (C) FRET efficiency of siCT and siCCM2 on 1 or 10kPa gel compared per category: linear (blue), ellipsoid (green), small (yellow) and large circular (purple) FAs. Statistical analysis between all groups was performed using linear-mixed effects model on the whole dataset (see Methods). N = 80, 80, 61, 60 for siCT-1kPa, siCT-10kPa, siCCM2-1kPa, siCCM2-10kPa respectively from 5 or 4 biological replicates, $*p < 0.05$, $**p < 0.01$, $***p < 0.001$.

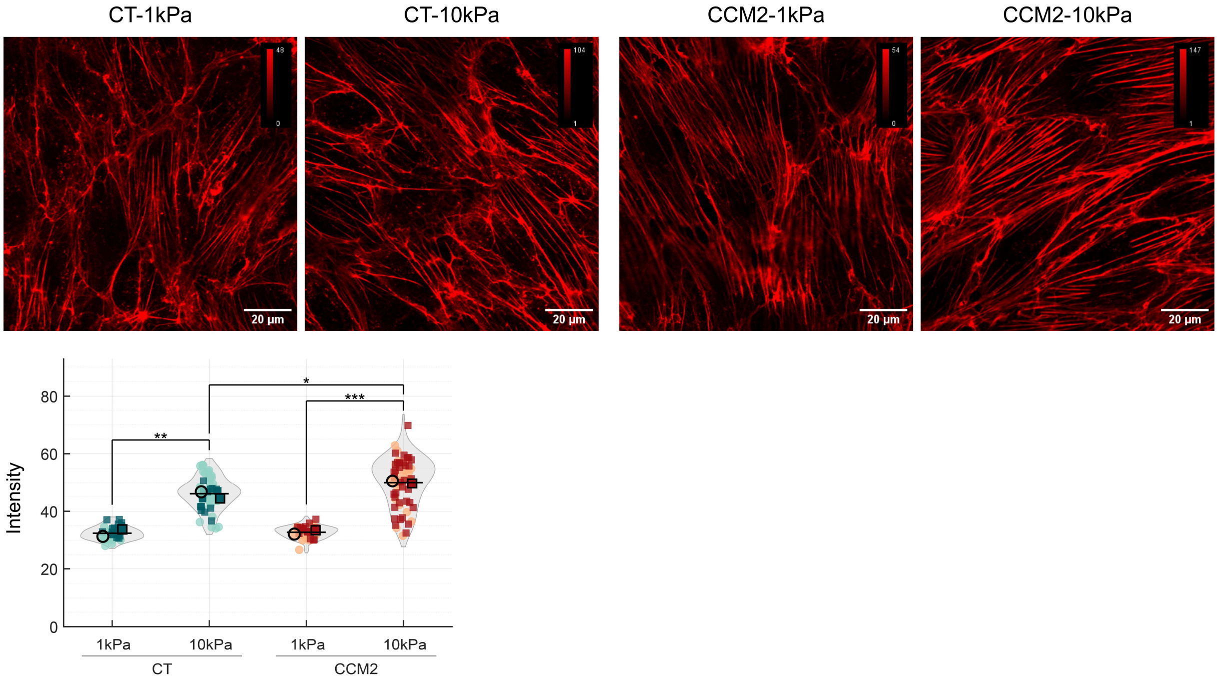


**Supplementary Figure 21: Actin staining in CT and CCM2 on 1kPa and 10kPa gel.** Top: SiR-actin stained HBMVEC monolayers show higher intensity for samples on 10kPa compared to 1kPa. Bottom: Average intensity of siCT and siCCM2 on 1 or 10kPa gel. Statistical analysis between all groups was performed using linear-mixed effects model (see Methods). N = 35, 56, 34, 62 for siCT-1kPa, siCT-10kPa, siCCM2-1kPa, siCCM2-10kPa respectively from 2 biological replicates, *$p < 0.05$, **$p < 0.01$, ***$p < 0.001$.

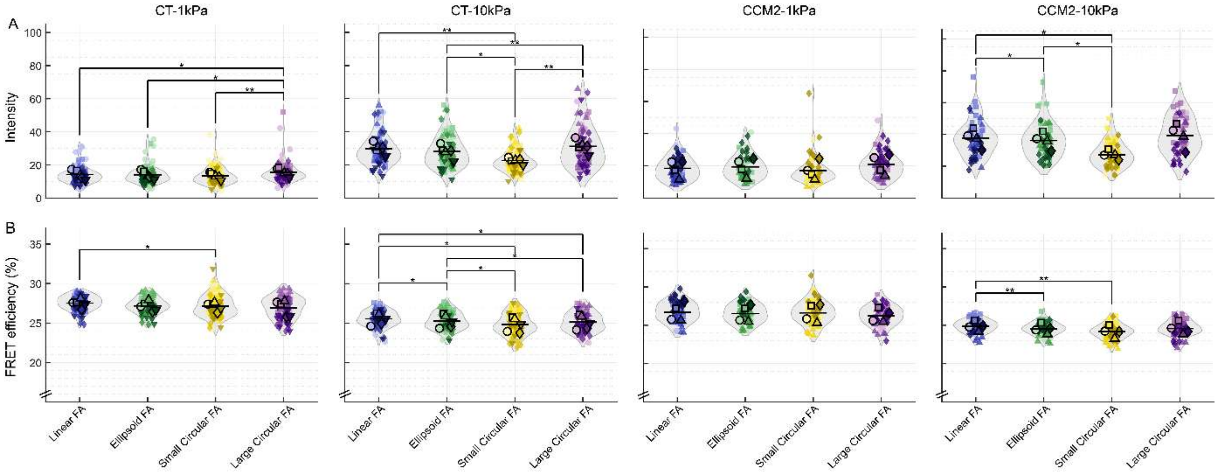


**Supplementary Figure 22: Comparison between morphological FA categories separated for CT and CCM2 on two gel stiffnesses.** The average (A) intensity and (B) FRET efficiency of per siCT-1kPa, siCT-10kPa, siCCM2-1kPa and siCCM2-10kPa condition per FA category. N = 80, 80, 61, 60 for siCT-1kPa, siCT-10kPa, siCCM2-1kPa, siCCM2-10kPa respectively from 5 or 4 biological replicates, *p < 0.05, **p < 0.01, ***p < 0.001.

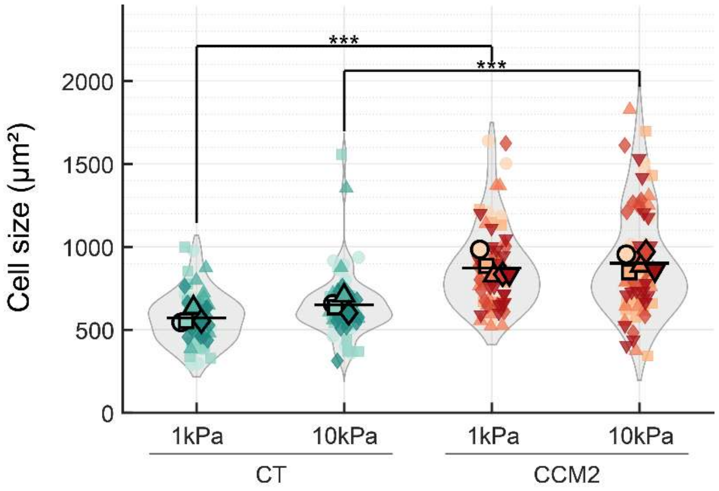


**Supplementary Figure 23: Comparison average cell size between CT and CCM2 on 1kPa and 10kPa gel.** N = 60, 60, 76, 79 for siCT-1kPa, siCT-10kPa, siCCM2-1kPa, siCCM2-10kPa respectively from 4 or 5 biological replicates, $*p < 0.05$, $**p < 0.01$, $***p < 0.001$.

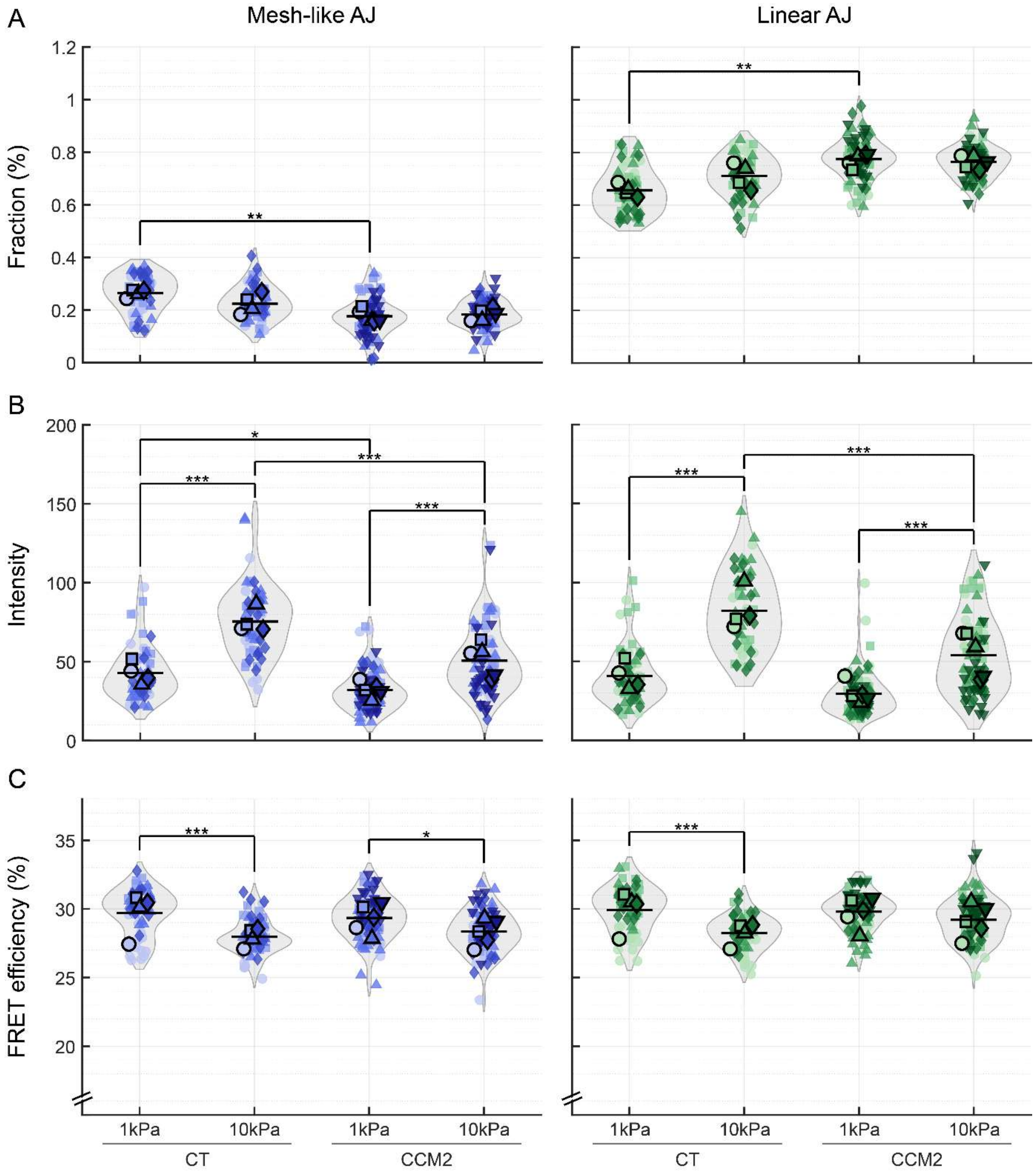


**Supplementary Figure 24: Comparison of CT and CCM2 on gel per morphological AJ category.** The average (A) fraction, (B) intensity and (C) FRET efficiency of siCT and siCCM2 on 1 or 10kPa gel compared per category: mesh-like (blue) and linear (green) AJs. Statistical analysis between all groups was performed using linear-mixed effects model on the whole dataset (see Methods). N = 60, 60, 76, 79 for siCT-1kPa, siCT-10kPa, siCCM2-1kPa, siCCM2-10kPa respectively from 4 or 5 biological replicates, $*p < 0.05$, $**p < 0.01$, $***p < 0.001$.

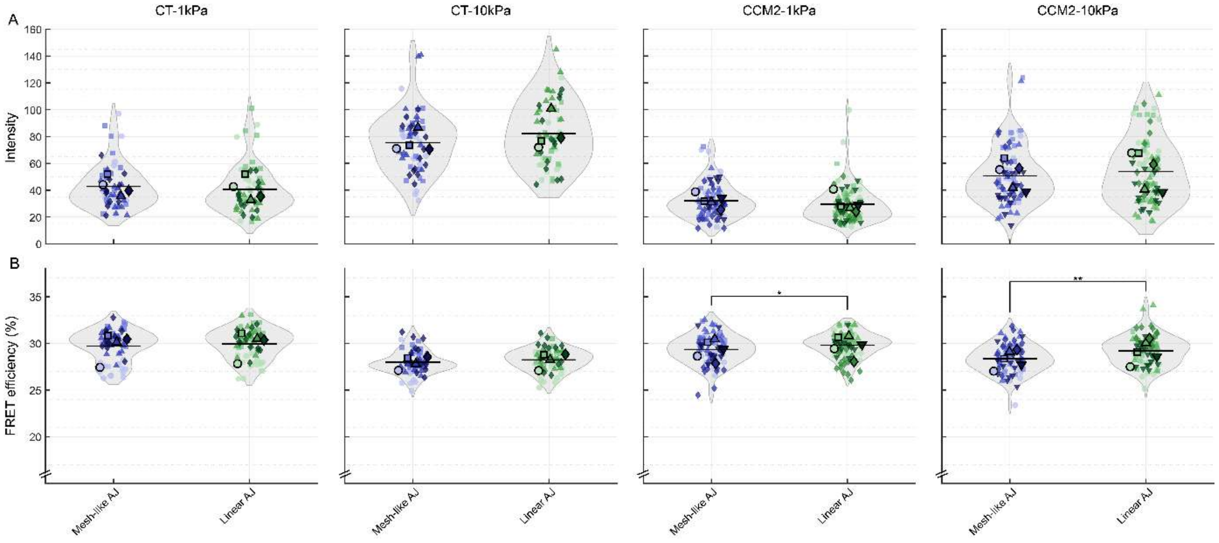


**Supplementary Figure 25: Comparison between morphological AJ categories separated for CT and CCM2 on two gel stiffnesses.** The average (A) intensity and (B) FRET efficiency of per siCT-1kPa, siCT-10kPa, siCCM2-1kPa and siCCM2-10kPa condition per AJ category. N = 60, 60, 76, 79 for siCT-1kPa, siCT-10kPa, siCCM2-1kPa, siCCM2-10kPa respectively from 4 or 5 biological replicates, *p < 0.05, **p < 0.01, ***p < 0.001.
