## Supplementary Notes for "Letting go to hold on: loss of CCM2 disrupts endothelial force balance in cerebral cavernous malformations"

### Supplementary note 1: Primers used

| Primer name | Sequence (5'-3') |
|---|---|
| **TSMods** | |
| TSMod – F40 | |
| Clover FW | TACGACTCACTATAGGGAGACCCAAGCTGGCTAGCGCCGCCACCATGG |
| Clover F40 REV | TCCCGGACCGGAACCGGCGGCGGTCACGAACTC |
| F40 (codon opt.) F | TTCGTGACCGCCGCCGGTTCCGGTCCGGGAGGTGCTGGC |
| F40 (codon opt.) REV | TCGCCCTTGGATCCAGCTCCA |
| mScarlet-I F40 FW | TGGAGGAGCCGGTCCAGGTGGAGCTGGATCCAAGGGCGAGGCAGTGATC |
| mScarlet-I REV | AGCGGCCGCCACTGTGCTGGATATCTGCAGAATTCTTACTTGTACAGCTCGTCCATG |
| TSMod – FL | |
| Clover FW | (see TSMod – F40 above) |
| Clover FL REV | GTCAAACTCGCCCATGGAACCGGCGGCGGTCACGAACTC |
| FL FW | ACCGCCGCCGGTTCCATGGGCGAGTTTGACATC |
| FL REV | CTCGCCCTTGGATCCTTCGAGGGAACCTCTCTC |
| mScarlet-I FL FW | AGAGGTTCCCTCGAAGGATCCAAGGGCGAGGCAGTGATC |
| mScarlet-I REV | (see TSMod – F40 above) |
| TSMod – HP35st | |
| Clover FW | (see TSMod – F40 above) |
| Clover HP35st REV | GTCCTCATCGCTCAGGGAACCGGCGGCGGTCACGAACTC |
| HP35st FW | ACCGCCGCCGGTTCCCTGAGCGATGAGGACTTC |
| HP35st REV | CTCGCCCTTGGATCCGAAGAGTCCCTTTTCCTTCATC |
| mScarlet-I HP35st FW | GAAAAGGGACTCTTCGGATCCAAGGGCGAGGCAGTGATC |
| mScarlet-I REV | (see TSMod – F40 above) |
| **pcDNA3.1-VECadTS** | |
| pcDNA3.1-VECadTL-F40 | |
| VECadherinHead FW | ACTCACTATAGGGAGACCCAAGCTGGCTAGCATGCAGAGGCTCACAGAGCT |
| VECadherinHead REV | TTGCTCACCATGGTAAGCTTACCGCGCACAGAATTAAGCA |
| TSMod FW | CTGTGCGCGGTAAGCTTACCATGGTGAGCAAGG |
| TSMod REV (pcDNA3.1 REV) | TAGAAGGCACAGTCGAGG |
| pcDNA3.1-VECadTS-F40 | |
| VECadherinTail1 FW | ACTCCACCGGCGGCATGGACGAGCTGTACAAGGCGGCC |
| VECadherinTail1 REV | GGCTTCTGCACCTGCGTATAGACTGCCGGCC |
| VECadherinTail2 FW | ACGCCCGGCCGGCAGTCTATACGCAGGTGCAG |
| VECadherinTail2 REV | CCGCCACTGTGCTGGATATCTGCAGAATTCCTAGATGATGAGTTCCTCCTGGGGATC |

### Supplementary note 2: Synthesized sequences

Codon optimized F40 sequence (5'-3')

GGTTCCGGTCCGGGAGGTGCTGGCCCCGGTGGTGCTGGTCCTGGTGGTGCAGGACCTGGTGGAGCA
GGGCCTGGAGGGGCTGGACCCGGGGGTGCGGGTCCTGGAGGAGCCGGTCCAGGTGGAGCTGGATC
CAAGGGCGAA

SFFV-EcoRI-NheI-WPRE-3'LTR cassette (5'-3')

GCAGAGATCCAGTTTGGATCGATAAGCTTGATATCGAATTTGCAGCCCCGATAAAATAAAAGATTTTATT
TAGTCTCCAGAAAAAGGGGGGAATGAAAGACCCCACCTGTAGGTTTGGCAAGCTACTGCAGTAACGC

CATTTTGCAAGGCATGGAAAAATACCAAACCAAGAATAGAGAAGTTCAGATCAAGGGCGGGTACATGA
AAATAGCTAACGTTGGGCCAAACAGGATATCTGCGGTGAGCAGTTTCGGCCCCGGCCCGGGGCCAAG
AACAGATGGTCACCGCAGTTTCGGCCCCGGCCCGAGGCCAAGAACAGATGGTCCCCAGATATGGCCC
AACCCTCAGCAGTTTCTTAAGACCCATCAGATGTTTCCAGGCTCCCCCAAGGACCTGAAATGACCCTGC
GCCTTATTTGAATTAACCAATCAGCCTGCTTCTCGCTTCTGTTCGCGCGCTTCTGCTTCCCGAGCTCTATA
AAAGAGCTCACAACCCCTCACTCGGCGCGCCAGTCCTCCGACAGACTGAGTCGCCCGGGGGGGATCT
GGAGCTCTCGAAATTCTCACGCGTCAAGTGGAGCAAGGCAGGTGGACAGTGGATCGCCGCTAGCTTG
CTCTGGGTCCTGCTGCTGAATTCGATCCTTTCAAGCTTATCGATAATCAACCTCTGGATTACAAAATTTGT
GAAAGATTGACTGGTATTCTTAACTATGTTGCTCCTTTTACGCTATGTGGATACGCTGCTTTAATGCCTTT
GTATCATGCTATTGCTTCCCGTATGGCTTTCATTTTCTCCTCCTTGTATAAATCCTGGTTGCTGTCTCTTTAT
GAGGAGTTGTGGCCCGTTGTCAGGCAACGTGGCGTGGTGTGCACTGTGTTTGCTGACGCAACCCCCA
CTGGTTGGGGCATTGCCACCACCTGTCAGCTCCTTTCCGGGACTTTCGCTTTCCCCCTCCCTATTGCCAC
GGCGGAACTCATCGCCGCCTGCCTTGCCCGCTGCTGGACAGGGGCTCGGCTGTTGGGCACTGACAAT
TCCGTGGTGTTGTCGGGGAAATCATCGTCCTTTCCTTGGCTGCTCGCCTGTGTTGCCACCTGGATTCTG
CGCGGGACGTCCTTCTGCTACGTCCCTTCGGCCCTCAATCCAGCGGACCTTCCTTCCCGCGGCCTGCTG
CCGGCTCTGCGGCCTCTTCCGCGTCTTCGCCTTCGCCCTCAGACGAGTCGGATCTCCCTTTGGGCCGCC
TCCCCGCATCGATACCGTCGACCTCGAAATTAATTCGAGCTCGGTACTTTAAGACCAATGACTTACAAGG
CAGCTGTAGATCTTAGCCACTTTTTAAAAGAAAAGGGGGGACTGGAAGGGCTAATTCACTCCCAACGA
AGACAAGATCTGCTTTTTGCTTGTACTGGGTCTCTCTGGTTAGACCAGATCTGAGCCTGGGAGCTCTCT
GGCTAACTAGGGAACCCACTGCTTAAGCCTCAATAAAGCTTGCCTTGAGTGCTTCAAGTAGTGTGTGCC
CGTCTGTTGTGTGACTCTGGTAACTAGAGATCCCTCAGACCCTTTTAGTCAGTGTGGAAAATCTCTAGCA
GCATCTAGAATTAATTCCGTGTATTCTATAGTGTCACCTAAATCGTATGTGTATGATACATAAGGTTATGTAT
TAATTGTAGCCGCGTTCTAACGACAATATGTAAAGCCTAATTGTGTAGCATCTGGCTTACTGAAGCAGAC
CCTATCATCCATCGCTCCAGATCCTCTACGCCGGACGCATCGTGGCCGGCA

### Supplementary note 3: qPCR primers

| Gene | Primer sequence | Source |
|---|---|---|
| CCM2 FW | CCTGCACAGCGATGACTCT | Human |
| CCM2 REV | ACCACCCACATCCACAGATT | |
| ROCK1 FW | AAAGAAAGGATGGAGGATGAAGT | Human |
| ROCK1 REV | TGTAACAACAGCCGCTTATTTG | |
| ROCK2 FW | CCCATCAACGTGGAGAGC | Human |
| ROCK2 REV | AAGCAGGAAAATCTAAATCAAGGA | |
